# Cross-species analysis of melanoma enhancer logic using deep learning

**DOI:** 10.1101/2019.12.21.885715

**Authors:** Liesbeth Minnoye, Ibrahim Ihsan Taskiran, David Mauduit, Maurizio Fazio, Linde Van Aerschot, Gert Hulsemans, Valerie Christiaens, Samira Makhzami, Monika Seltenhammer, Panagiotis Karras, Aline Primot, Edouard Cadieu, Ellen van Rooijen, Jean-Christophe Marine, Giorgia Egidy Maskos, Ghanem-Elias Ghanem, Leonard Zon, Jasper Wouters, Stein Aerts

## Abstract

Genomic enhancers form the central nodes of gene regulatory networks by harbouring combinations of transcription factor binding sites. Deciphering the combinatorial code by which these binding sites are assembled within enhancers is indispensable to understand their regulatory involvement in establishing a cell’s phenotype, especially within biological systems with dysregulated gene regulatory networks, such as melanoma. In order to unravel the enhancer logic of the two most common melanoma cell states, namely the melanocytic and mesenchymal-like state, we combined comparative epigenomics with machine learning. By profiling chromatin accessibility using ATAC-seq on a cohort of 27 melanoma cell lines across six different species, we demonstrate the conservation of the two main melanoma states and their underlying master regulators. To perform an in-depth analysis of the enhancer architecture, we trained a deep neural network, called DeepMEL, to classify melanoma enhancers not only in the human genome, but also in other species. DeepMEL revealed the presence, organisation and positional specificity of important transcription factor binding sites. Together, this extensive analysis of the melanoma enhancer code allowed us to propose the concept of a core regulatory complex binding to melanocytic enhancers, consisting of SOX10, TFAP2A, MITF and RUNX, and to disentangle their individual roles in regulating enhancer accessibility and activity.

## Introduction

A cell’s phenotype arises from the expression of a unique set of genes, which is regulated through the binding of transcription factors (TFs) to cis-regulatory elements, such as promoters and enhancers. Deciphering gene regulatory programs entails understanding the network of transcription factors and cis-regulatory elements that governs the identity of a given cell type; as well as understanding how the specificity of such a network is encoded in the DNA sequence of genomic enhancers. Enhancers harbor combinations of binding sites for TFs, through which transcription of nearby target genes is regulated^1,2^. The chromatin around enhancers is typically enriched for acetylation of histone H3 at lysine 27 (H3K27ac) and H3 monomethylation at K3 (H3K4me1), allowing enhancer identification through ChIP-seq for these specific histone marks^1^. In addition, profiling accessible chromatin via DNase I hypersensitive sequencing (DNase-seq) or via the Assay for Transposase-Accessible Chromatin using sequencing (ATAC-seq) represents a useful approach for identifying putative enhancers^3,4^. Indeed, active enhancers are typically depleted of one or more nucleosomes, due to the binding of TFs. Initial changes in DNA accessibility can be facilitated through a special class of TFs that bind with high affinity to their recognition sites and that have a long residence time at the enhancer; sometimes referred to as pioneer TFs^4,5^. By displacing nucleosomes or thermodynamically outcompeting nucleosome binding they allow other TFs to co-bind, thereby further stabilising the nucleosome depleted region and/or actively enhancing transcription of target genes^6,7^. As the presence and architecture of TF binding sites within enhancers determine which TFs can bind with high affinity, understanding this ‘enhancer logic’ can help interpreting the functional role of enhancers within a gene regulatory network. Several techniques exist to study the enhancer code, including (1) motif discovery tools, in which position-weight matrices of TF binding sites are used to calculate their enrichment in sets of co-regulated regions or co-expressed genes^8,9^; (2) comparative genomics, by exploiting cross-species data to identify conserved and therefore possible important (parts of) enhancers^10–12^; (3) genetic screens to measure the effect of mutations on enhancer activity^13,14^; and (4) machine learning techniques, where mathematical models are trained to recognise specific patterns in enhancers and help to classify them^15^. Particularly the latter has seen a strong boost the last years, with the advent of large training sets derived from genome-wide profiling. Three pivotal methods based on deep learning include DeepBind^16^, DeepSEA^17^ and Basset^18^, the first convolutional neural networks (CNNs) applied to genomics data^19^. Since their emergence in the genomics field, machine learning techniques, and especially CNNs, have been applied to model a range of regulatory aspects, including TF binding sites^20^, DNA methylation^21^ and 3D chromatin architecture^22^, by exploiting large epigenomics datasets.

Deciphering gene regulation and the underlying enhancer code is not only important during dynamic processes such as development, but also in disease contexts such as cancer, where gene regulatory networks are typically dysregulated due to mutations. Melanoma is a type of skin cancer which mostly develops from a buildup of UV-induced mutations in melanocytes, the pigment-producing cells in the skin^23^. Particularly in this cancer type, gene expression is dysregulated and highly plastic, giving rise to two main melanoma cell states: the melanocytic (MEL) state, which still resembles the cell-of-origin, i.e. the melanocyte, expressing high levels of the melanocyte-lineage specific transcription factors MITF, SOX10 and TFAP2, as well as typical pigmentation genes such as *DCT, TYR, PMEL*, and *MLANA*; and the mesenchymal-like (MES) state, in which the cells are more invasive and therapy resistant, expressing low levels of MITF and SOX10, and high levels of genes involved in TGFbeta signaling and epithelial-to-mesenchymal transition (EMT)-related genes^24–28^. These transcriptomic differences have also been studied at the epigenomics level, with AP-1 and TEAD factors as master regulators of the MES state and binding sites for SOX10 and MITF significantly enriched in MEL-specific regulatory regions^27–29^. However, it remains unclear how these regulatory states are encoded in particular enhancer architectures, and whether such architectures are evolutionary conserved. Besides human cell lines and human patient-derived cultures, several animal models have been established in melanoma research, including mouse, pig, horse, dog and zebrafish^30–34^. Although these models are widely used, it is unknown whether their enhancer landscapes and regulatory programs are conserved with human.

Here, we combine comparative regulatory genomics with machine learning to investigate enhancer logic in melanoma. Through epigenomic profiling of 27 melanoma cell lines across six species, we examine the conservation of the two main melanoma states and underlying master regulators. By training a deep neural network, called DeepMEL, on topic models derived from the human cell lines, we were able to classify not only human melanoma enhancers, but also regulatory regions in the other species. DeepMEL revealed high-confidence TF binding sites for the different melanoma states, how they are positioned within melanoma enhancers, and where they are placed with respect to the central enhancer nucleosome. This in-depth analysis of the melanoma enhancer code allowed us to propose a mechanistic model of TF binding in MEL melanoma enhancers. Finally, by exploiting the deep layers of our model, we are able to identify causal mutations for melanoma enhancer loss and gain through evolution, not only affecting enhancer accessibility but also activity.

## Results

### Melanoma chromatin accessibility landscapes are conserved across species

To study the conservation of melanoma cell states and underlying enhancer logic, we performed (Omni)ATAC-seq on a cohort of melanoma cell lines across six species, obtaining accessible chromatin landscapes of a total of 27 samples (Fig. 1a). These include 17 human patient-derived cultures (“MM lines”)^27,35^, one mouse cell line^36^, one cell line derived from the pig melanoma model MeLiM (“MeLiM”)^30^, two horse melanoma lines derived from a Grey Lipizzaner horse (“HoMel-L1”) and from an Arabian horse (“HoMel-A1”)^33^, two dog melanoma cell lines (“Cesar” and “Bounty”)^37^ and four melanoma lines established from zebrafish (“ZMEL1”, “EGFP-121-1”, “EGFP-121-5” and “EGFP-121-3”)^38,39^. Per sample, between 65,475 and 176,695 ATAC-seq peaks were observed (Fig. S1a), including regions that are accessible across all six species in this study and thus conserved (e.g. *TCF7L2* promoter), peaks that are only accessible in the mammalian lines (i.e. in human, mouse, pig, horse and dog lines) (e.g. *ST3GAL2* promoter) and species-specific peaks (e.g. the human-specific *NMNAT1* intronic enhancer) (Fig. 1a). Interestingly, unsupervised clustering of the 17 human lines grouped the samples into two distinct clusters (Fig. S1b), which correspond to the two main cell states in human melanoma, i.e. the melanocytic state (MEL) and mesenchymal-like state (MES), as was confirmed for twelve of the lines by RNA-seq data using established MEL and MES gene signatures (Fig. S1c)^27^. Indeed, regulatory regions near MEL-specific genes such as *SOX10* were accessible in human lines in the MEL state (MM001, MM011, MM031, MM034, MM052, MM057, MM074, MM087, MM118, MM122 and MM164), whereas they were closed in MES melanoma lines (MM029, MM047, MM099, MM116, MM163, and MM165) (Fig. 1b). In addition, this classification was in agreement with previous work were respectively nine and ten of these lines were clustered based on epigenomic data (using OmniATAC-seq, and H3K27ac ChIP-seq and FAIRE-seq, respectively)^27,28^. Of note, similarly as in Wouters et al., we observed inter-cell line heterogeneity within the states, especially within the melanocytic state (Fig. S1b).

To examine whether the two main melanoma states were conserved in the other species of our cohort, we first identified conserved regulatory regions using the liftOver tool^40^ to compare genomic positions. Between 1.1% and 40.9% of the ATAC-seq regions in non-human lines were conserved in human, i.e. convertible to human coordinates and accessible in human; and between 0.9% and 18.4% of the human peaks were conserved in the other species (Fig. 1c). Note that the most distant species in our cohort, i.e. zebrafish (last common ancestor ∼340 million years ago^41^), has the smallest proportion of conserved regions (1.1%), as expected. Accordingly, we identified 10,592 regulatory regions conserved across the mammalian species, and, when including zebrafish, 116 conserved regions across all six studied species (Fig. 1d). Nearly half of the 10,592 conserved mammalian regions were promoters within 1 kb of a transcription start site (Fig. 1d). Indeed, high conservation of proximal promoters has previously been reported, which is partially due to their position near the transcription units, making them evolutionarily more stable compared to more distal regulatory elements^12^. In each of the mammalian species, the 10,592 conserved regions were more accessible compared to all ATAC-seq regions and, in addition, these conserved regions show a higher ChIP-seq signal for H3K27ac in human, a mark for active regulatory regions^42^ (Fig. S1d,e).

**Figure 1.**
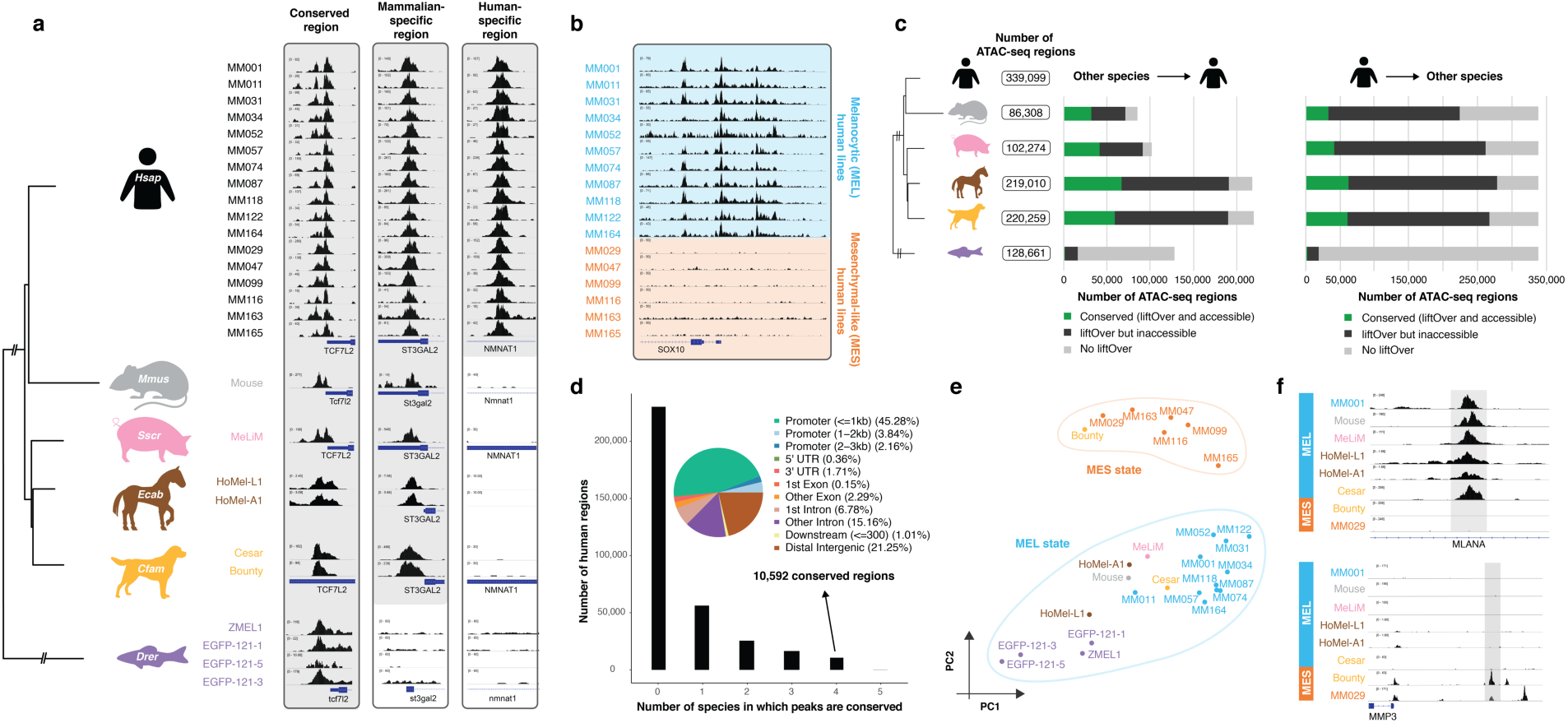
Comparative epigenomics reveals conservation of two main melanoma states. **a**, Evolutionary relationship between the six studied species, represented by a phylogenetic tree (NCBI taxonomy tree). ATAC-seq profiles of the 27 melanoma cell lines are shown for a conserved region (*TCF7L2* promoter), a mammalian-specific region (*ST3GAL2* promoter) and a human-specific region (*NMNAT1* intronic enhancer). **b**, ATAC-seq profiles of the human melanoma lines for the *SOX10* locus. Lines are coloured by the melanocytic (MEL, in blue) or mesenchymal-like (MES, in orange) melanoma state. **c**, (left) Total number of ATAC-seq regions observed across all samples of a species, (middle) coloured based on their liftOver (at least 10% of bases must remap) and conservation status compared to human. (right) Similar graph for the conservation of the 339,099 human regions in each of the other species. **d**, Number of human regions that are conserved with 0 (i.e. human-specific) to 5 different species. ChIPseeker results are shown for the 10,592 human regions that are conserved across all mammalian species. **e**, Melanoma cell lines cluster into two groups, linked to the MEL and MES melanoma states as shown in a PCA plot based on 116 conserved regions across all six species. **f**, ATAC-seq profiles of MEL and MES lines of different species for an intronic *MLANA* enhancer and the upstream region of *MMP3*.

Next, to test how closely related the different melanoma lines are at the epigenomic level, we clustered the lines using the identified conserved regions. Clustering of all mammalian samples based on the accessibility of the 10,592 conserved mammalian regions (Fig. S1g,h) or of all samples using the 116 globally conserved regions (Fig. 1e, Fig. S1f), revealed again two main clusters. One cluster contained all human MEL samples together with 9 of the 10 non-human lines, indicating that most of the non-human cell lines are epigenomically similar to human MEL lines. On the other hand, the second cluster consisted of all human MES samples together with the dog cell line ‘Bounty’. Based on this co-clustering of melanoma lines, we can state that all non-human cell lines are in the MEL state, except for the dog line ‘Bounty’ which belongs to the MES state. Indeed, known MEL regulatory regions such as the intronic enhancer of *MLANA*, a MEL-specific gene involved in melanosome biogenesis^43^, are accessible in all mammalian lines, except for the MES human lines and the dog line Bounty; whereas the opposite is true for an enhancer upstream of *MMP3*, a gene which increases metastatic potential in melanoma cell lines^44^ (Fig. 1f).

In conclusion, by using ATAC-seq on a panel of 27 melanoma lines across six species, conserved regulatory regions could be identified. These regions allowed clustering of the melanoma samples into two groups which correspond to the two main melanoma cell states, indicating conservation of the MES melanoma state in dog and the MEL melanoma state in pig, mouse, horse, dog and even zebrafish melanoma samples.

### Conserved transcription factor motifs determine state-specific enhancers

Next, we wanted to investigate whether the conserved MEL and MES states are controlled by similar master regulators across different species. First, we performed an evolutionary comparison of differential transcription factor binding sites between MEL and MES cell lines in human and dog, as these were the only two species in our cohort for which cell lines of both states were available. Differential peak calling between the human MEL and MES lines revealed significant enrichment of SOX, TFAP2, MITF, RUNX and ETS TF binding motifs in the 25,164 differential MEL human peaks (log2FC > 2.5 and pAdj < 0.0005; complete Homer output in Supplementary Table 1) (Fig. 2a). Indeed, SOX10, TFAP2 and MITF are among the previously reported master regulators of the MEL state^24,27–29^. The 12,824 human differential MES regions were significantly enriched for binding motifs for transcription factors of the AP-1 family and TEADs (Fig. 2a), known regulators in human MES melanoma lines^27^. To examine the conservation of these master regulators in dog melanoma, we contrasted the two dog lines. Interestingly, the 58,515 peaks specific to the MEL dog line Cesar were significantly enriched for similar TF binding motifs as the human differential MEL peaks, i.e. SOX, TFAP2, RUNX, MITF and ETS motifs, and even the order of the enriched TF families was comparable (Fig. 2b). The same was true for the motifs enriched in the MES-specific human and dog regions (Fig. 2b). Note that the difference in the number of differentially accessible regions between dog and human is likely due to the variability between human samples that are used as replicates, while for dog we used two technical replicates of the same cell line. Altogether, these observations indicate that the MEL and MES melanoma cell states are conserved in dog and that they are likely governed by the same master regulators, based on the concordance of motif enrichment for SOX10, MITF, TFAP2 and ETS factors; and for AP-1 and TEAD TFs for the MEL and MES state respectively.

**Figure 2.**
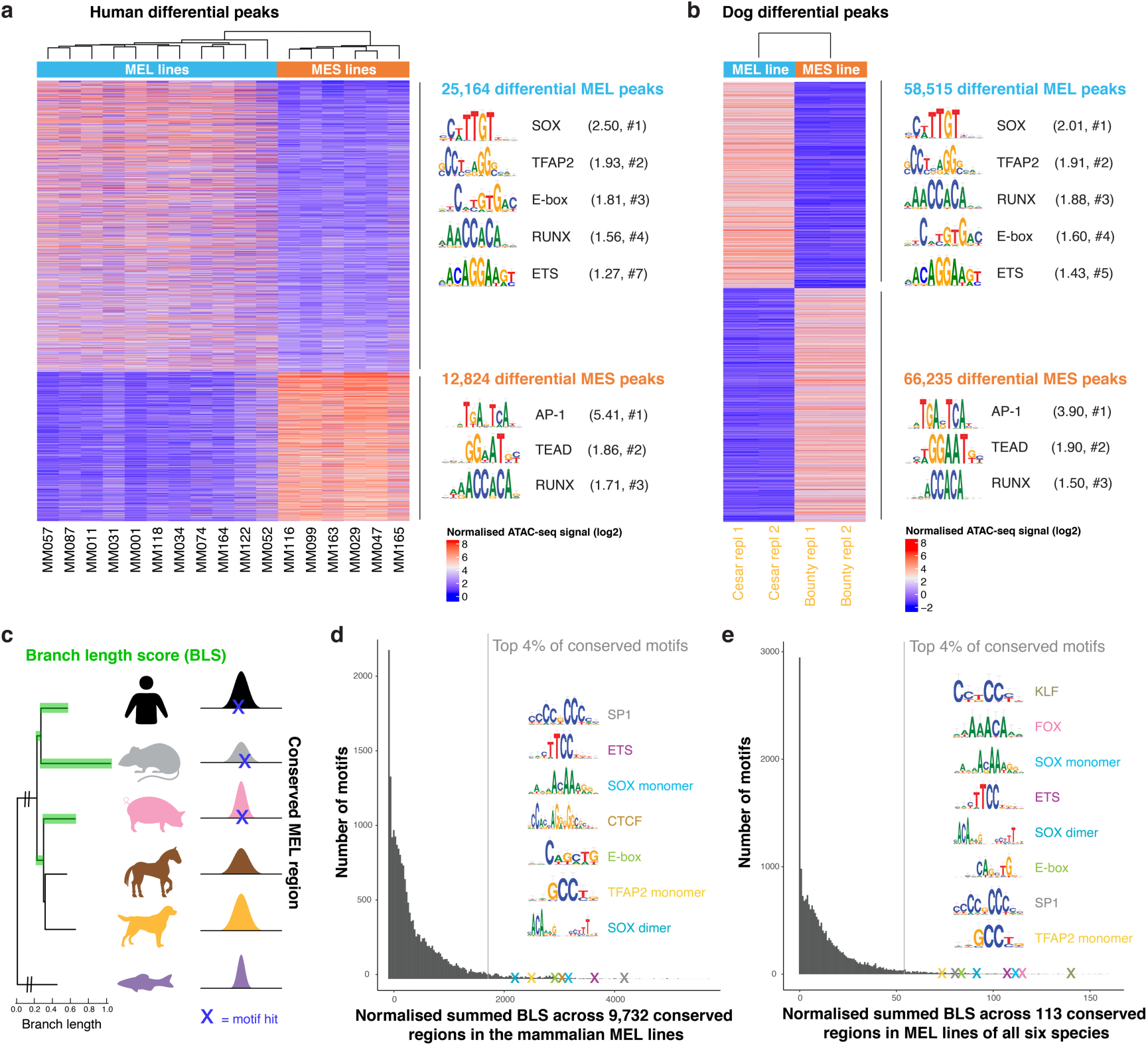
Conservation of binding motifs of master regulators of MEL and MES melanoma states. **a, b**, Heatmap of differential ATAC-seq regions when comparing (**a**) human MEL versus human MES lines and (**b**) the MEL dog line ‘Cesar’ versus the MES dog line ‘Bounty’ (two biological replicates each), coloured by normalised ATAC-seq signal. Enriched TF binding motifs in the differential peaks were identified via Homer^47^ and the first logo of enriched TF families is shown. The ratio of the percentage of target sequences with the motif and the percentage of background sequences with the motif is indicated between brackets, as well as the rank of the TF class within the Homer output (#). **c**, Schematic overview of cross-species motif analysis using the branch length score (BLS) as a measure for the evolutionary conservation of a motif hit (for 20,003 TF position-weight matrices) across conserved regions. The BLS was summed across a set of conserved regions, i.e. the higher the BLS score, the more conserved the motif is in that specific set of regions. **d, e**, Histogram of the normalised summed BLS score for 20,003 motifs on (**d**) 9,732 conserved regions across the mammalian MEL lines and on (**e**) 113 conserved regions across MEL lines of all six species. The first hit of the top recurrent TF binding motifs within the top 4% conserved motifs is indicated as a cross and is accompanied by the logo of the motif.

To further verify the importance of the MEL-specific master regulators in MEL cell lines of the remaining four species, we applied a different strategy since we could not contrast MEL and MES lines for horse, pig, mouse and zebrafish. Therefore, we focused on 9,732 regions that were conserved across all mammalian MEL lines to identify conserved TF binding sites. Note that this number differs from the 10,592 conserved regions defined above as only the MEL lines were used here. We scanned the 9,732 conserved regions using our library of 20,003 TF position-weight matrices (PWMs) and used a branch length score (BLS) to calculate the level of evolutionary conservation of each TF binding motif (Fig. 2c), a strategy applied before in other systems^7,45^. Among the 4% most conserved motifs were SP1, ETS, SOX (both monomer and dimer motifs), CTCF, MITF and TFAP2 motifs (Fig. 2d). Notably, the top conserved motifs were members of the SP/KLF TF family, which bind to GC-rich motifs in promoters^46^. Indeed, 47% of the 9,732 conserved regions in mammalian MEL lines were proximal promoters (<= 1 kbp from TSS). BLS scoring on the remaining 5,196 more distal conserved regions showed no longer conservation SP1/KLF TF motifs, but just conservation of the previously identified TF binding motifs for TFAP2A, MITF, SOX10, CTCF and ETS factors (Fig. S1i), indicating that distal regions, such as enhancers, mostly contain the state-specific TF binding motifs. Interestingly, when we included zebrafish ATAC-seq regions, only 113 regions were conserved in the MEL cell lines across all six species, but BLS scoring still revealed SOX, ETS, MITF and TFAP2 motifs among the most conserved motifs in MEL lines (Fig. 2e). Note that we did not perform any contrast of MEL versus MES lines prior to the BLS analyses and that these motifs were identified by just focusing on the conserved regions in MEL melanoma lines.

Altogether, two independent strategies of motif analysis suggest that melanoma enhancer logic is conserved across species and that the MEL state is governed by conserved master regulators including SOX10, MITF, TFAP2A and ETS.

### Deep neural network DeepMEL reveals nucleotide-resolution enhancer logic

While motif enrichment can predict candidate regulators, we sought to build a more comprehensive model of the MEL enhancers, that would allow cross-species predictions and in-depth analysis of enhancer architecture. To this end, we trained a deep learning (DL) model on human ATAC-seq data. First, to construct an unsupervised training set, we clustered all 339,099 human ATAC-seq peaks using cisTopic^48^ (see Methods) into 24 topics (Fig. 3a, Fig. S2a,b). This provided a more nuanced classification, with topic 4 representing the MEL enhancers being accessible across all MEL samples; and topic 7 representing the MES enhancers that are accessible in the MES samples (Fig. 3a, Fig. S2c). In addition, we found two topics containing regions that are generally accessible across all cell lines (topic 1 and topic 19) (Fig. 3a, S2c), and which were highly enriched for proximal promoters (Fig. S2d) and for known promoter-specific TF binding motifs linked to SP1 and NFY TF families (Fig. S2c)^46,49^. Other topics were more specific to one or a small group of cell lines. For instance, topic 22 contained regions that were mostly accessible in MM057, MM074 and MM087 (Fig. 3a). These particular lines have previously been reported as an ‘intermediate’ (INT) sub-state of the MEL state, governed by a mixed MEL-MES GRN^28^. We verified the biological relevance of these topics by investigating nearby target genes using GREAT^50^. Genes near topic 4 regions are significantly enriched for Gene Ontology (GO) terms such as pigmentation (FDR=1.95e-8) and neural crest cell differentiation (FDR=4.26e-7), whereas genes near topic 7 regions were more mesenchymal-like as they are enriched for GO terms involved in cell-cell adhesion (1.56e-13). Next, we performed motif discovery on the top regions assigned to each topic. SOX, ETS, TFAP2 and MITF motifs were enriched in regions of the MEL-specific topic 4 and AP-1 in the MES-specific topic 7 (Fig. S2c), confirming our findings from the supervised differential peak calling discussed above (Fig. 2a). An example topic 4 region in the promoter of the SOX10 target gene *MIA*^51^ is shown in Figure 3b, as well as two topic 7 regions upstream of *SERPINE1*, a gene expressed in metastatic melanoma^52^.

**Figure 3.**
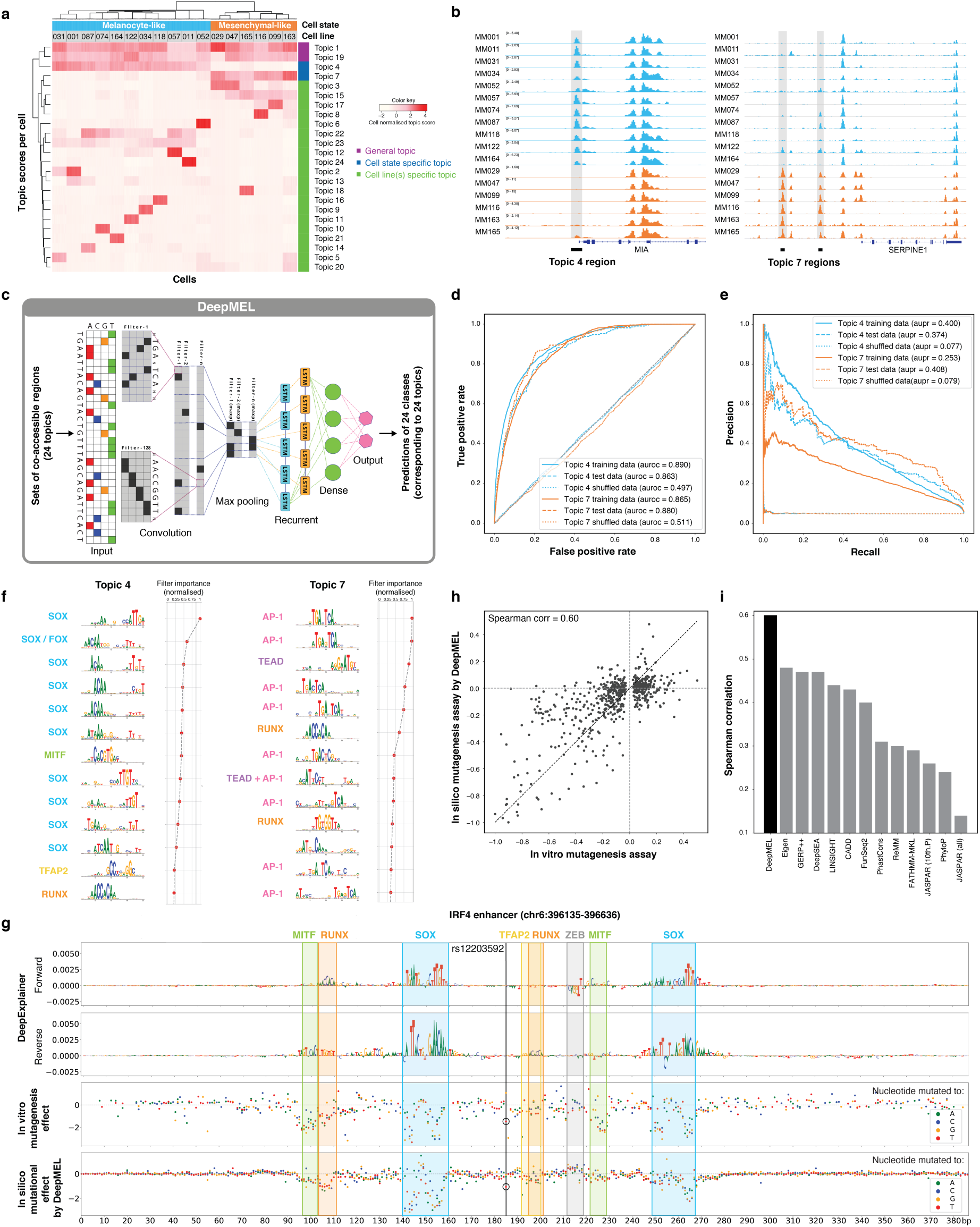
DeepMEL classifies melanoma enhancers and predicts important TF binding motifs. **a**, Cell-topic heatmap of cisTopic applied to 339,099 ATAC-seq regions across the 17 human melanoma lines, coloured by normalised topic scores. 24 topics or sets of co-accessible regions are found, including general topics, cell state specific topics and cell line(s) specific topics. **b**, Example regions of a MEL-specific (topic 4) region near *MIA* and MES-specific (topic 7) regions upstream of *SERPINE1*. **c**, Schematic overview of DeepMEL. 24 sets of co-accessible regions were used as input for training of a multi-class multi-label neural network. **d, e**, (**d**) Receiver operating characteristic curve and (**e**) precision-recall curve for DeepMEL on training, test and shuffled data of topic 4 and topic 7 regions. **f**, Top 13 enriched filters learned by DeepMEL to classify regions as MEL (topic 4) or MES (topic 7). Filters were converted to logos and accompanied by the candidate TF binding motif names, as identified by TomTom comparison^59^. Normalised filter importance is shown per filter. **g**, Example of a MEL-predicted enhancer near *IRF4*. (first and second row) DeepExplainer view of the forward and reverse strand are shown, with the height of the nucleotides indicating the importance for prediction of the MEL enhancer. SOX, MITF, TFAP2, ZEB-like and RUNX-like motifs within the enhancer are highlighted. (third row) *In vitro* effect of point mutations on enhancer activity as measured by MPRA^14^. Colours represent the nucleotide to which the wild type nucleotide is mutated. (bottom row) *In silico* effect of point mutations as predicted by DeepMEL. The location of SNP rs12203592 is highlighted by a black vertical line and the *in vitro* and *in silico* point mutations that generate the SNP are encircled. **h**, Correlation between the *in vitro* mutational effects on the *IRF4* enhancer compared to the *in silico* mutagenesis predictions. **i**, Performance of variant effect prediction of several previously tested models on the *IRF4* enhancer case^14^.

Using the 24 topics as classes, we trained a multi-class, multi-label classifier using a neural network, called “DeepMEL” (Fig. 3c). As input, we used the forward and reverse complement of 500 bp enhancer sequences centered on the ATAC-seq summit. As topology, we used the DanQ CNN-RNN hybrid architecture^53^ consisting of 4 main layers: a convolution layer to discover local patterns in sequential data, followed by a max-pooling layer to reduce the dimensionality of the data and generalise the model effectively, a bidirectional recurrent layer (LSTM) to detect long-range dependencies of the local patterns discovered in the first layer, and finally a fully-connected (dense) layer just before the output layer to help the classification after the feature extraction layers (Fig. 3c). After successful training of DeepMEL (auroc = 0.863 and aupr = 0.374 on test data for topic 4 regions) (Fig. 3d,e, Fig. S3a), we used the weights of neurons from the convolutional filters to extract local patterns learned by the model. We transformed these convolution filters into PWMs and found the importance of each filter for each topic (see Methods and Supplement). Intriguingly, filters that represent SOX, MITF, TFAP, and RUNX motifs were most relevant for the MEL-specific topic 4 and filters that represent AP-1, TEAD and RUNX binding sites were assigned to the MES-specific topic 7 (Fig. 3f). Thus, DeepMEL learned the relevant features *de novo* from the sequence. DeepMEL can be used to score and classify any given DNA sequence of 500 bp. For instance, when re-entering all ATAC-seq peaks of the MEL line MM001 in the model, it classified 3,885 regions as MEL-specific (topic 4 scores above threshold of 0.16 (see Methods)). These regions were indeed highly accessible in MEL lines and closed in MES lines, and interestingly, were also accessible in human melanocytes (Fig. S3b,c). Importantly, this indicates that these MEL-specific regions in melanoma are not cancer-specific but already accessible in their cell-of-origin, i.e. the melanocytes, and that we potentially can extrapolate the observations on this topic to melanocyte enhancers. Although in the remainder of this work we will score accessible regions to identify functional enhancers, it is also possible to score the entire genome, without filtering for ATAC-seq peaks. This may be useful for species where no ATAC-seq data of melanoma or melanocytes is available. Such a scoring yields high precision and recall (69% and 86% respectively, Fig. S3d).

In order to examine the TF binding site architecture within enhancers, we used a model interpretation tool, DeepExplainer^54,55^, which does backpropagation of the activation differences^56^, to visualise the importance of each nucleotide in an enhancer with respect to the predicted enhancer class. For instance, in a MEL enhancer located on the 4th intron of *IRF4*, nucleotides important for classifying this enhancer as topic 4 form motifs for SOX10, MITF, TFAP and RUNX factors (Fig. 3g top two rows). Indeed, SOX10 binding has been reported on this location^57^. Another example is given for a region of topic 22, the topic specific to the INT MEL subpopulation, where SOX10 and AP-1 co-exist within the same enhancer, indicating that these cell lines also contain properties of a mixture between the MEL and MES state at the epigenomic level (Fig. S3e,f).

Importantly, it is known that enhancer accessibility does not directly translate to enhancer activity^1^. To test whether the same TF binding motifs were contributing to the activity of MEL enhancers, we used the *IRF4* enhancer as case study. For this enhancer, Kircher et al.^14^ performed saturation mutagenesis followed by an *in vitro* massively parallel reporter assay (MPRA), testing the effect of every possible single nucleotide mutation on enhancer activity (Fig. 3g, 3th row). The most deleterious mutations coincided with the SOX, E-box and RUNX-like motifs that were predicted by DeepMEL, indicating that the predicted motifs are also contributing to enhancer activity, as their disruption reduced enhancer activity *in vitro*. To further examine how well DeepMEL can predict the *in vitro* MPRA effect, we measured the effect on the topic 4 DL score of each single nucleotide mutation *in silico* (Fig. 3g, bottom row). Interestingly, mutations that have the strongest *in silico* effect overlapped with predicted TF binding motifs, and more intriguingly, also the magnitude of the effect highly correlated with the *in vitro* mutations (Spearman correlation of 0.60) (Fig. 3g,h), even though DeepMEL was trained only on binary accessibility data (i.e. binary topics of co-accessible regions). These observations indicate that, although the DeepMEL was trained to predict enhancer accessibility, it is also a good predictor of enhancer activity of this specific enhancer. Notably, our DeepMEL performed best in predicting the *in vitro* mutagenesis on the *IRF4* enhancer activity compared to other classifiers and deep learning models that were benchmarked in Kircher et al.^14^ (CAGI challenge, 2018) (Fig. 3i). Interestingly, enhancer accessibility and activity were not only influenced by mutations that break a motif for an activating TF, but also by the creation of a repressor binding motif. This was the case for a C-to-T mutation that coincided with a SNP involved in freckles, brown hair and high sensitivity of the skin to sun exposure (rs12203592, SNPedia) (Fig. 3g). This SNP creates a ZEB/SNAI-like motif that negatively contributes to the MEL topic score of this enhancer (Fig. S3g). A similar motif was also found to decrease the MEL prediction in the wild-type sequence (Fig. 3g, “ZEB”, letters facing downwards) and mutating this motif increased the topic 4 prediction score, indicating that the ZEB/SNAI-like TF binding motif (CAGGT) may function as a repressor for the MEL state. Indeed, ZEB factors have been reported to act as transcriptional repressors by interaction with the corepressor CtBP^58^, and mutations in the binding motif of the transcriptional repressor SNAI2 have been shown to increase chromatin accessibility^11^. Note that the ability of DeepMEL to predict the effect of mutations on enhancer accessibility (and activity) raises the opportunity to apply DeepMEL to predict enhancer mutations that affect chromatin accessibility in, for instance, personalised cancer genomes; as we did in our companion paper for phased melanoma genomes of a total of 10 patient-derived melanoma cultures (Kalender Atak et al., 2019).

In conclusion, our DL model DeepMEL, trained on topics of human co-accessible regions, is performant in classifying melanoma regulatory regions into different classes based on purely the DNA sequence. Interestingly, features learned by DeepMEL corresponded to TF binding motifs of master regulators of specific classes. These motifs could also be located and visualised within regions using a model interpretation tool, allowing examination of the motif architecture within specific enhancers and predicting the effect of mutations on enhancer accessibility.

### Cross-species scoring identifies orthologous melanoma enhancers

Next, we wanted to use the human-trained DL model DeepMEL for predicting MEL and MES enhancers in other species. We started with the dog genome as a test case, because the differential ATAC-seq peaks between the MEL (Cesar) and MES (Bounty) dog cell lines could be used as true positives. DeepMEL reached an area under the receiver operating characteristic (auroc) of 0.979 for predicting MEL regions (as topic 4) versus MES regions (as topic 7) in dog, which approximates the model’s performance for classifying human MEL and MES differential regions (auroc = 0.987), and this accuracy is significantly higher compared to using cis-regulatory module (CRM) scoring with PWMs (Fig 4a,b,c). Having confirmed that the human model can identify enhancers in the dog epigenome, we predicted MEL and MES enhancers across all six species. This yielded between 2,093 and 5,400 MEL enhancers, and between 7,459 and 10,743 MES enhancers, in samples of the MEL and MES state respectively (Fig. 4d, S4c). Interestingly, although the total number of accessible regions in the genome varies between cell lines and species (Fig. 4d, numbers between brackets), for all MEL cell lines around 2.5% of the accessible regions were predicted MEL enhancers. Note that the majority of these enhancers could not have been detected using whole genome alignments (liftOver) (Fig. 4b,c, Fig S4a-d).

**Figure 4.**
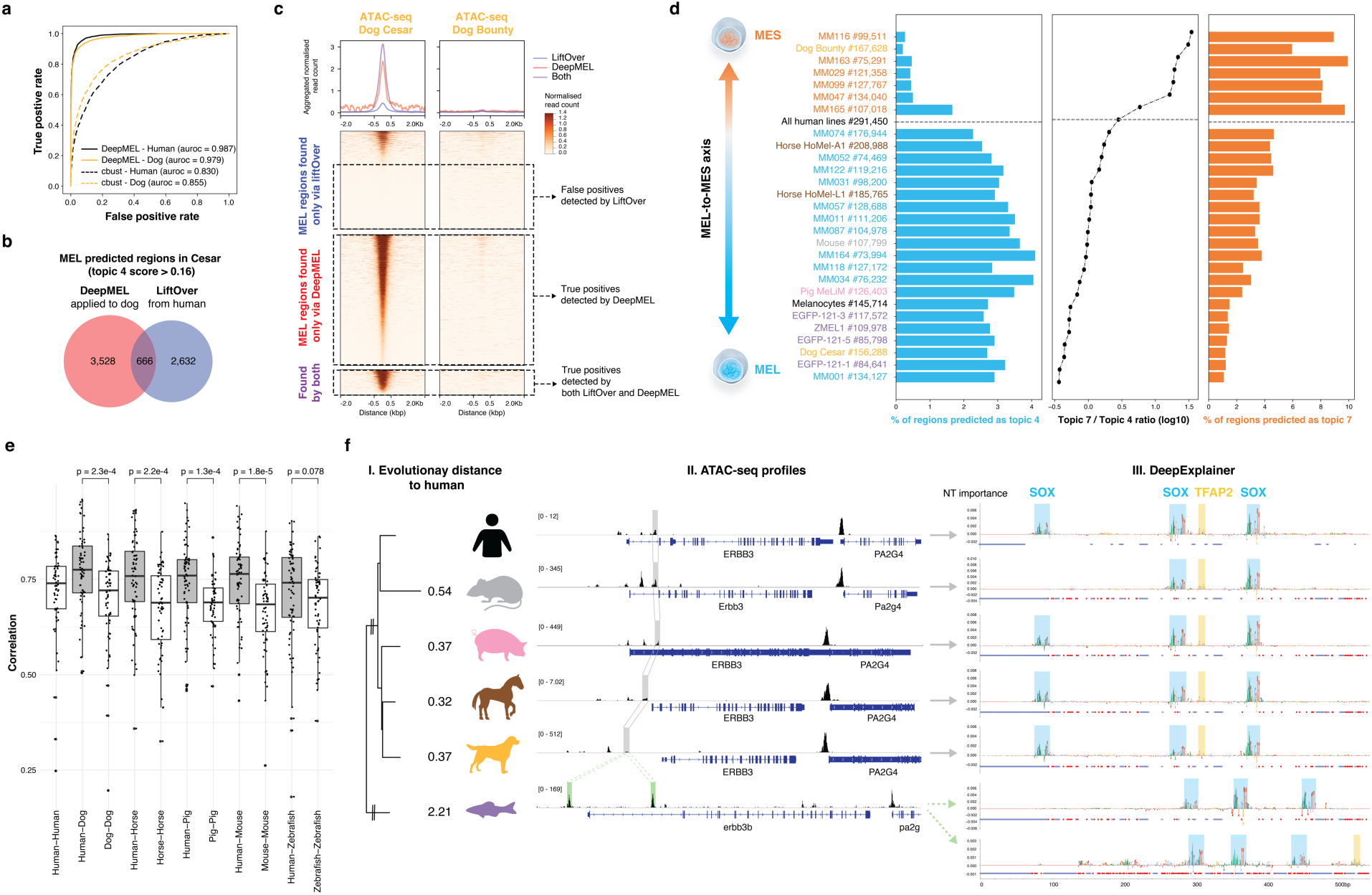
Human-trained deep learning model on cross-species ATAC-seq data. **a**, DeepMEL performs well in classifying MEL and MES differential peaks in human and dog, and outcompetes Cluster-Buster (cbust). **b**, Venn diagram of the number of topic 4 (MEL-specific) regions predicted by DeepMEL in the dog line ‘Cesar’ and of dog regions found by liftOver of the human MEL regions. **c**, Heatmaps of ATAC-seq signal of the dog lines ‘Cesar’ and ‘Bounty’ on MEL-predicted regions found via liftOver (blue), MEL regions predicted by DeepMEL (red) and MEL regions identified by both methods (purple). Heatmaps are coloured by normalised read counts and ordered according to the ATAC-seq signal in ‘Cesar’. Aggregation plots are shown on top. **d**, Percentage of MEL and MES predicted ATAC-seq regions across all samples in our cohort and in human melanocytes. Samples are ordered according to the MEL-MES axis by using the ratio of the number of MES / MEL predicted regions. **e**, Pearson correlation of deep layer scores between MEL-predicted regions of orthologous MEL genes between human and another species (‘Human-Species’) or between MEL-predicted regions near different MEL genes within one species (‘Species-Species’). **f**, (I) Evolutionary distance between human and other species in branch length units. (II) ATAC-seq profiles of the *ERBB3* locus in the six different species. MEL-specific enhancers that were predicted by DeepMEL and that were also found via liftOver of the human MEL enhancer are highlighted in grey, whereas MEL-predicted regions only found by DeepMEL are highlighted in green. (III) DeepExplainer plots are shown for the multiple-aligned MEL-predicted *ERBB3* enhancers, for zebrafish the first and second row represent the DeepExplainer plots of the upstream and intronic enhancer, respectively. SOX and TFAP2 motifs formed by important nucleotides are highlighted. Red and blue dots represent point and indels mutations, respectively.

Having identified high-confidence MEL enhancers genome-wide across 6 species, as a combination of ATAC-seq peaks and high topic 4 prediction scores, we analysed their distribution with respect to orthologous genes, and their evolutionary divergence. Particularly, we looked at enhancers located near a set of 379 human genes that are specifically expressed in the MEL state (derived from RNA-seq data across a cohort of twelve MM lines (see Methods)). Of these 379 genes, 217 (67%) had at least one MEL-predicted enhancer within a locus of 200kb up- and downstream of the gene (the MEL cell line MM001 was used for this analysis). Between 70-85% of the orthologous MEL genes in other species had at least one MEL enhancer nearby (Fig. S4e). Note that only a small subset of these enhancers could have been found using liftOver (2-43% depending on the species). Of these genes, 32 form a core set of conserved genes throughout all species, each having a MEL enhancer, including zebrafish. Examples of genes in the core set are *MITF, PMEL* and *TYRP1*, genes known to be involved in melanocyte development, melanosome formation and melanin production^60^.

A long-standing question in enhancer studies is how to compare enhancers with each other, if their sequences do not align^61,62^. Here we tackle this question by using the dense layer of DeepMEL as a reduced dimensional space to calculate the correlation between enhancers. Using this measure we found that MEL-predicted enhancers in proximity of homologous MEL genes are significantly more similar to each other compared to MEL-predicted enhancers in proximity of different MEL genes within the same species (Fig. 4e), indicating that MEL enhancers near orthologous genes are indeed orthologous enhancers. Note that the correlation of orthologous MEL enhancers approximated or even surpassed the correlation of redundant (or shadow enhancers^63^) linked to the same MEL gene in a species (Fig. S4f). Lastly, we studied an example of a MEL enhancer in more detail, namely the enhancer near *ERBB3*. DeepMEL predicts a MEL enhancer upstream or intronic of *ERBB3* in each of the mammalian species, which were also found by liftOver of the human *ERBB3* enhancer (Fig. 4f II). However, in the zebrafish genome, liftOver was unable to identify the homologous region, whereas DeepMEL predicted two MEL enhancers, one upstream of the TSS of *erbb3b* and another in the first intron. Both zebrafish enhancers were highly correlated with the human ERBB3 enhancer (deep layer pearson correlation of 0.812 and 0.797 for the upstream and intronic zebrafish enhancer, respectively), suggesting that both enhancers are orthologous to the human *ERBB3* enhancer. Applying DeepExplainer to the multiple-aligned sequences revealed a conserved motif architecture in the orthologous mammalian *ERBB3* enhancers containing each three SOX motifs and one TFAP2 motif (Fig. 4f III). Note that in mouse, one SOX binding site was lost, mouse is also the mammalian species that is most distant from human, among the included species in this study (Fig. 4f I). The two zebrafish enhancers contain several SOX motifs, however with different inter-motif distances. The two zebrafish enhancers have a highly similar motif architecture, suggesting that they arose by duplication from a common ancestor enhancer.

In conclusion, we showed that DeepMEL is able to identify MEL- and MES-specific enhancers in different species, which allows studying evolutionary events and enhancer logic within orthologous enhancers, even in distant species such as zebrafish.

### Motif architecture of the MEL enhancer

To study the architecture of MEL enhancers in more detail, including motif composition, motif order and distance, and relationships to the nucleosome position, we set out to obtain high-confidence motif annotations in each of the 3,885 MEL enhancers in human (MM001, the most MEL-like human cell line), for each of the predicted core regulatory factors (SOX10, MITF, TFAP2A, RUNX). To achieve this, we devised an improved motif scoring method that obtains precise positions of TF binding motifs by multiplying DeepMEL activation scores of convolutional filters (i.e. motifs) with the DeepExplainer profile on each enhancer (Fig. 5a)^64^. A motif hit is predicted as significant when its importance is above a motif-specific threshold which was determined by using all regions as background (see Methods).

**Figure 5.**
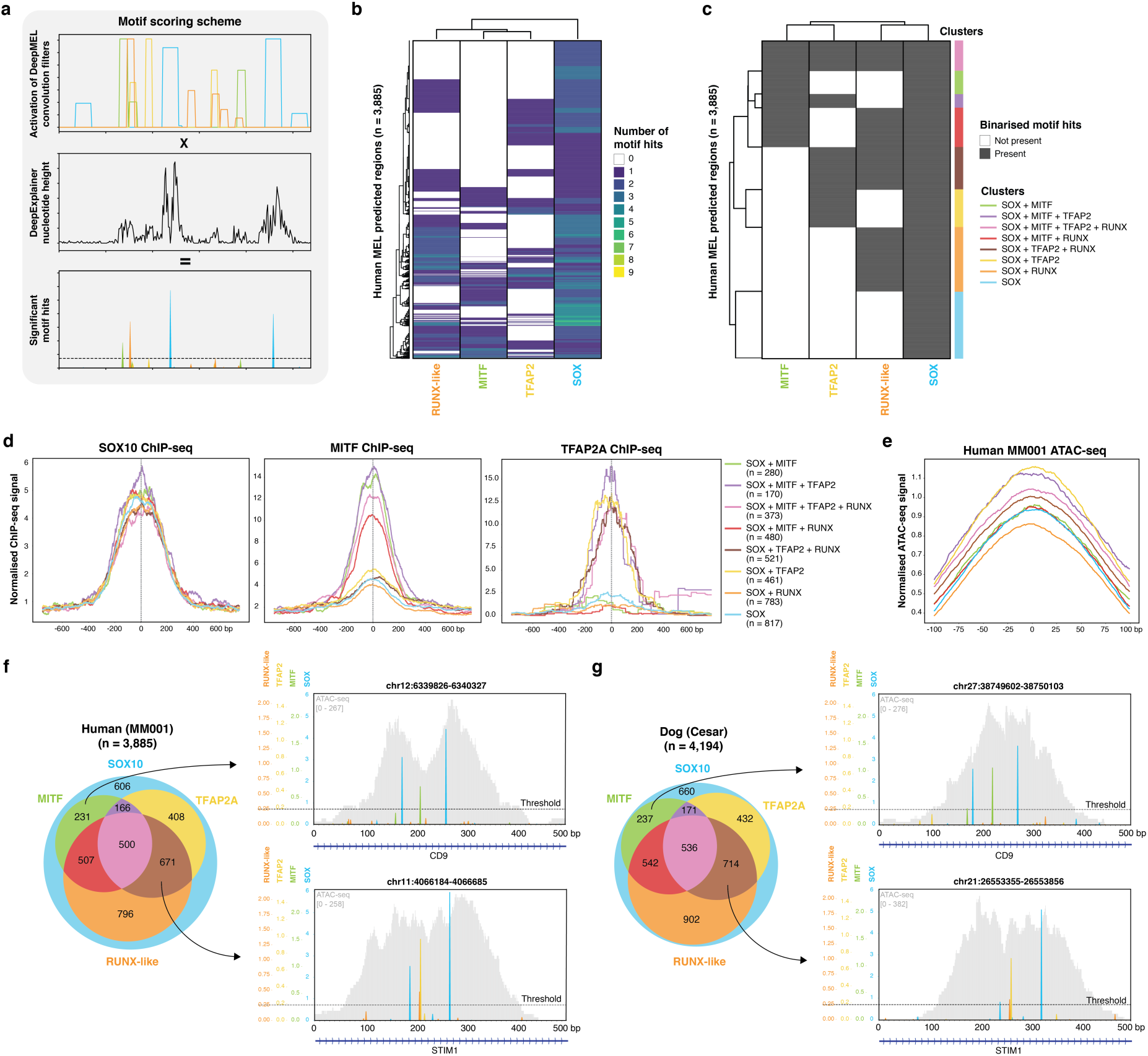
COre Regulatory Complex of MEL melanoma enhancers. **a**, Schematic overview of motif scoring method in which extended convolutional filter hits from DeepMEL are multiplied by DeepExplainer profiles to yield significant motif hits. **b**, Heatmap of the number of significant SOX, TFAP2, MITF and RUNX-like motif hits on the 3,885 MEL predicted regions in the human cell line MM001. **c**, Binarised heatmap of significant SOX, TFAP2, MITF and RUNX-like motif hit(s) on the 3,885 MEL predicted regions in the human cell line MM001. Eight region clusters can be distinguished, representing different combinations of significant motifs present in the enhancers. **d**, Aggregation plot of normalised ChIP-seq signal of SOX10 (left), MITF (middle) and TFAP2A (right) on the human enhancer clusters. **e**, Aggregation plot of normalised ATAC-seq signal of MM001 on the human enhancer clusters. **f, g**, Venn diagram representation of regions clusters on (**f**) the 3,885 regions predicted as MEL in human (in MM001) and (**g**) the 4,194 MEL-predicted regions in dog (in Cesar). In addition, example MEL-predicted regions in human and dog are shown for two of the region clusters: an intronic *CD9* enhancer as representative for the SOX10 + MITF cluster and an intronic *STIM1* enhancer containing SOX10, TFAP2 and RUNX motif hits.

The first remarkable observation was that each MEL enhancer contains at least one SOX10 motif hit, and often two or more (Fig 5b). This suggests that SOX10 plays a central role in MEL enhancer accessibility. Indeed, knock-down of SOX10 in MM001 significantly decreases the accessibility of MEL enhancers (Fig. S5a), and the regions that close after SOX10-KD are highly enriched for SOX motifs (NES = 28.5), possibly revealing a pioneering-role of SOX10 in MEL enhancers. Pioneer factors can access their binding sites on nucleosomal DNA, thereby directly or indirectly displacing the nucleosome, which results in the accessibility of the region^5^. Next to SOX, a combination of one or multiple TFAP2, MITF or RUNX-like motifs was present in 84% of the MEL-predicted enhancers. To facilitate a systematic study of the MEL enhancer logic, we binarised the motif-region matrix to simplify the region clustering (Fig 5c). We obtained 8 different enhancer classes, each with a different motif composition (Fig. 5c). As validation of the clusters and the predicted TF binding sites, we used human ChIP-seq data of SOX10, MITF and TFAP2A in melanoma or melanocytes^65,66^ (Fig. 5d). All clusters were indeed highly bound by SOX10, validating the prevalent SOX10 motif in all MEL enhancers. In contrast, MITF ChIP-seq data revealed that MITF binds more to enhancer classes with MITF motifs compared to regions lacking a significant MITF motif. Similarly, only enhancers containing at least one TFAP2 motif were bound by TFAP2A. Interestingly, regions containing a TFAP2A motif, next to the SOX10 motif(s) and possible others, showed a modest increase in accessibility (Fig. 5e), which could be in line with the previously described role of TFAP2A as a stabiliser of nucleosome-depleted regions^6^. The opposite was true for regions containing RUNX-like TF binding sites, as these were found to be less accessible compared to regions containing only SOX10 motifs, suggesting a repressive role of RUNX factors. The presence of a MITF site did not seem to alter the accessibility of enhancers compared SOX-only enhancers, but did increase H3K27ac signal (Fig. S5b), possibly indicating that MEL enhancers bound by MITF are more active.

To validate these MEL enhancer classes in other species, we applied the same motif scoring and binarisation to DeepMEL-predicted MEL regions in the other species in our cohort. Interestingly, MEL enhancers in other species also clustered into the same 8 clusters, with a similar distribution of regions per cluster (Fig. 5f,g, Fig. S5c). To test the conservation of the clusters, we used liftOver to compare the classification of enhancers across species. Although identifying orthologous sequences via whole genome alignment is not always correct, as shown above, a general trend was observed where the regions of a human cluster correspond to the same cluster in the other species (Fig. S5d), indicating conservation of the MEL enhancer clusters across species. For instance, the dog-orthologs of two human MEL enhancers belonging to either the cluster containing SOX10 and MITF binding sites (intronic enhancer of *CD9*) or to the cluster containing SOX10, TFAP2A and RUNX-like motifs (intronic enhancer of *STIM1*) (Fig. 5f) were part of the corresponding clusters in dog (Fig. 5g). In these examples we observed preserved spacing of around 80 bp between the two SOX10 binding sites within the enhancers, to which we will come back further below.

Altogether, these data suggest a COre Regulatory Complex (CoRC)^67^ of SOX10, TFAP2A, MITF and RUNX factors in regulating melanoma MEL enhancers, encoded by a mixed enhancer model^68^, with high flexibility in the combination of binding sites for these four TFs, but with some rigidity (or hierarchy) in the code as at least one SOX10 binding site is required.

### SOX10 functions as pioneer and TFAP2A as stabiliser in melanoma MEL enhancers

As previous results suggested a pioneering and stabiliser function for SOX10 and TFAP2A respectively, we wanted to further investigate these putative roles and how they are mechanistically affecting chromatin accessibility. First, we analysed the location of binding sites relative to the position of the nucleosome, focusing on MEL enhancers that contain a combination of SOX10 and TFAP2A sites (Fig. 6a,b). We predicted the nucleosome start and middle point using a previously published model^69^. Interestingly, we observed that SOX10 binding sites are situated within the borders of the nucleosome, near the former nucleosome start point, whereas TFAP2A binding occurs preferentially near the center of the nucleosome (Fig. 6a,b). Note that KD of TFAP2A halved the accessibility of this specific human region, whereas SOX10-KD completely abolished the ATAC-seq peak (Fig. 6a), indicating that SOX10 is necessary for accessibility, and that TFAP2A further increases the accessibility, which is in line with our previous observations (Fig. 5e, S5a).

**Figure 6.**
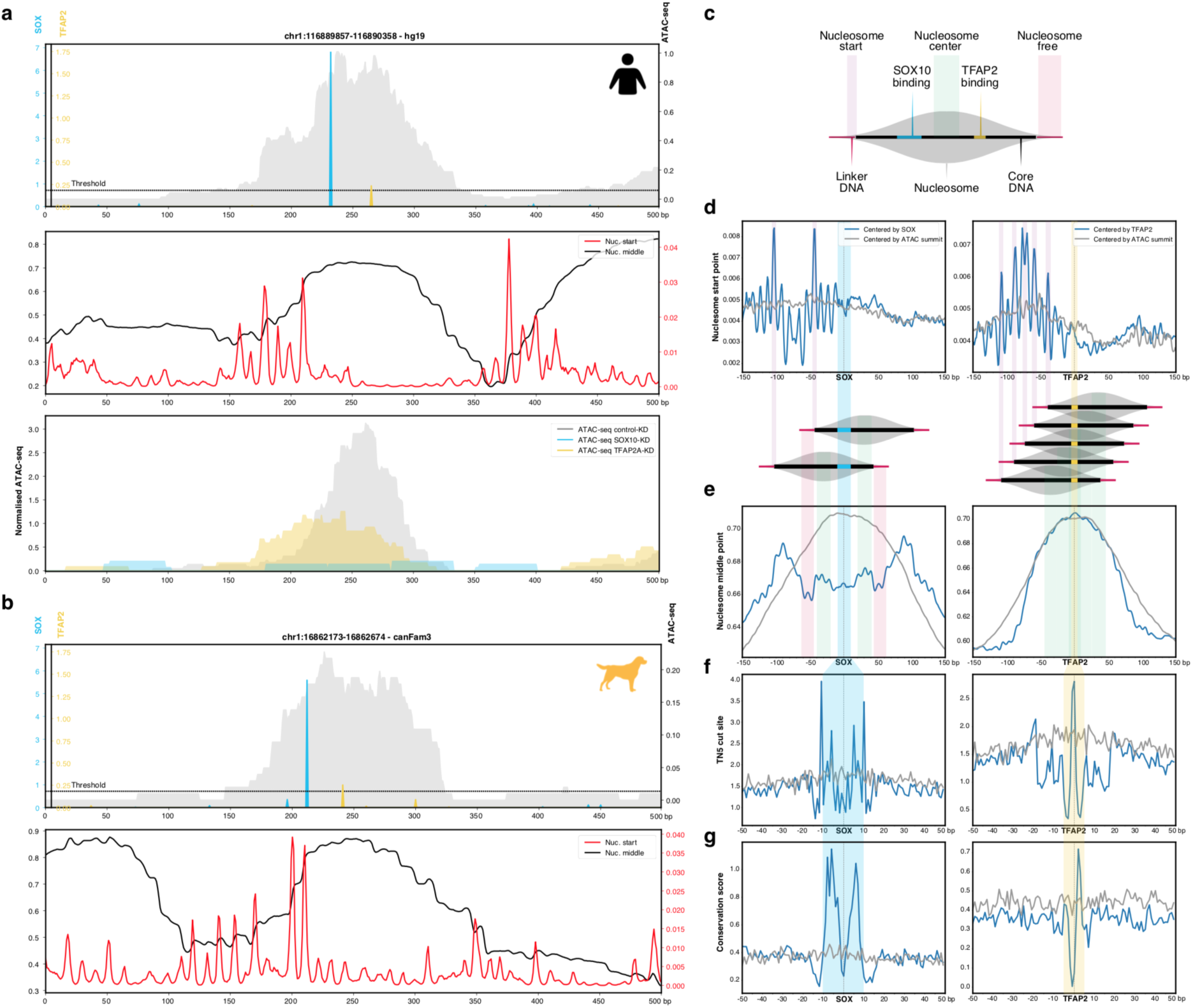
Positional specificity of SOX10 and TFAP2A in MEL melanoma enhancers. **a**. (top) Example human MEL-predicted enhancer containing significant SOX10 and TFAP2 motifs. The ATAC-seq signal is shown in grey. (middle) Imputed nucleosome start and middle point profiles. (bottom) ATAC-seq profiles of MM001 in control condition, after 72 h of SOX10 knock-down or TFAP2A knock-down. **b**. (top) Example dog MEL-predicted enhancer containing significant SOX and TFAP2 motifs. The ATAC-seq peak is shown in grey. (bottom) Imputed nucleosome start and middle point profiles. **c**. Schematic overview of nucleosome structure explaining the colours used in (**d**,**e**,**f**,**g**). **d**,**e**,**f**,**g**. Nucleosome start point (**d**), nucleosome middle point (**e**), Tn5 cut site (**f**), phyloP conservation score profiles (**g**) on MEL-predicted regions containing one SOX10 (left) or one TFAP2 motif (right) next to possible other motifs, where the regions are either centered on the ATAC-seq summit (grey) or on the SOX10 or TFAP2 motif (blue). SOX10 binding is enriched around 40 bp away the nucleosome start point, as is clear by the two peaks in the nucleosome start profile (**d**) that are situated respectively ∼40 and ∼110 bp away from the beginning of the SOX10 motif (which is 20 bp long), reflecting SOX binding at either side of the nucleosome as shown by the illustration.

These example enhancers raised an interesting positional preference of SOX10 and TFAP2A. To assess whether this occurs globally we centered human MEL enhancers on the SOX10 and TFAP2A motif hits and calculated the aggregated location of the nucleosome start and middle point (Fig. 6c,d,e). Interestingly, SOX10 had a consistent preference for binding within the nucleosome borders, around 40 bp away from the nucleosome start point (Fig. 6c,d). Since in chromatinised DNA, 146 bp of DNA sequence is wrapped around the nucleosome, we anticipated the nucleosome middle point to be situated ∼35 bp (= 146 bp / 2 - 40 bp) away from the SOX10 motif, which was indeed the case (Fig. 6e). Other pioneering factors have also been shown to bind near the borders of the nucleosome, such as FOX factors which bind around 60 bp from the center of the nucleosome, displacing linker histones and destabilising the central nucleosome^6,70^. On the other hand, when centering the MEL regions based on the TFAP2A motif, we did not observe a strong preference in the location of the nucleosome start point relative to the TFAP2A binding site (Fig. 6d), but in fact TFAP2A was consistently binding in a wide range on and around the nucleosome middle point (Fig. 6e). Stabilisators, such as NFIb, are known to directly compete with the central nucleosomes to stabilise the accessible chromatin configuration^6,71^. Centering based on the SOX10 motif hit revealed protection of Tn5 cutting on the conserved nucleotides of the dimer (Fig 6f,g). Similarly, protection and conservation was observed on important nucleotides in the TFAP2A dimer. We did not observe strong positional preferences of MITF and RUNX motifs relative to the nucleosome start or middle point (Fig. S6).

Altogether these data highly suggest that SOX10 functions as a pioneer in the CoRC of MEL enhancers, leading to their accessibility by binding to the central nucleosome, near the nucleosome start point. On the other hand, TFAP2A appears to act as stabiliser of SOX-dependent nucleosome depleted regions by binding around the nucleosome middle point, possibly going in competition with the central nucleosome.

### DeepMEL predicts evolutionary changes in MEL enhancer accessibility and activity

Next, we wanted to further validate our findings on the MEL enhancer logic using comparative genomics. This allowed us, in addition, to test how turnover of TF binding sites affects enhancer accessibility and function. To this end, we compared pairs of MEL enhancers that are homologous but only accessible in one of the species, to investigate which mutations cause the collapse of a MEL enhancer during evolution (Fig. 7a). We focused only on pairs of highly probable orthologous enhancers by requesting a stringent liftOver score (minimum of 99% of the bases must remap) and high sequence identity (at least 80% of the bases must be identical). We calculated the loss in ATAC-seq signal and in DeepMEL score, and aligned the sequence pairs to determine point mutations and indels between the homologous sequences (Fig. 7a). For example, an enhancer upstream of *APPL2* is predicted as MEL enhancer in the MEL dog line Cesar (topic 4 DL score of 0.35), whereas the orthologous enhancer in human was completely closed (Fig. 7b). Interestingly, not only the accessibility of the human homolog was lost, but also the activity, as we confirmed by a luciferase assay (Fig. 7c). Importantly, the DeepMEL score for this enhancer was seven times lower in human than in dog, falling below the topic 4 significance threshold of 0.16, indicating that the model detected critical changes in the human enhancer sequence that could explain the loss of this MEL enhancer. To determine which mutations were causal for the loss in accessibility (and activity), we calculated the effect on the MEL prediction score of each detected point mutation between the dog and human sequence, via *in silico* mutating the dog sequence (see Methods, similar as in the *IRF4* enhancer above). Several mutations seemed to alter the DL score (Fig. 7e,f). To pinpoint the functional effect of each mutation, we plotted DeepExplainer profiles and significant motif hits for CoRC factors on the original dog and human sequence (Fig. 7f). The functional dog enhancer contained a SOX10, MITF and TFAP2A binding site, which (almost) disappeared in the non-functional human homologous sequence. The losses could be explained by one T-to-A mutation in the SOX10 motif, one A-to-G mutation in the MITF motif and two mutations in the TFAP2A motif (Fig. 7f, encircled mutations). The SOX10 motif mutation had the strongest effect, as it caused a 45% drop in the MEL-prediction score (Fig. 7e).

**Figure 7.**
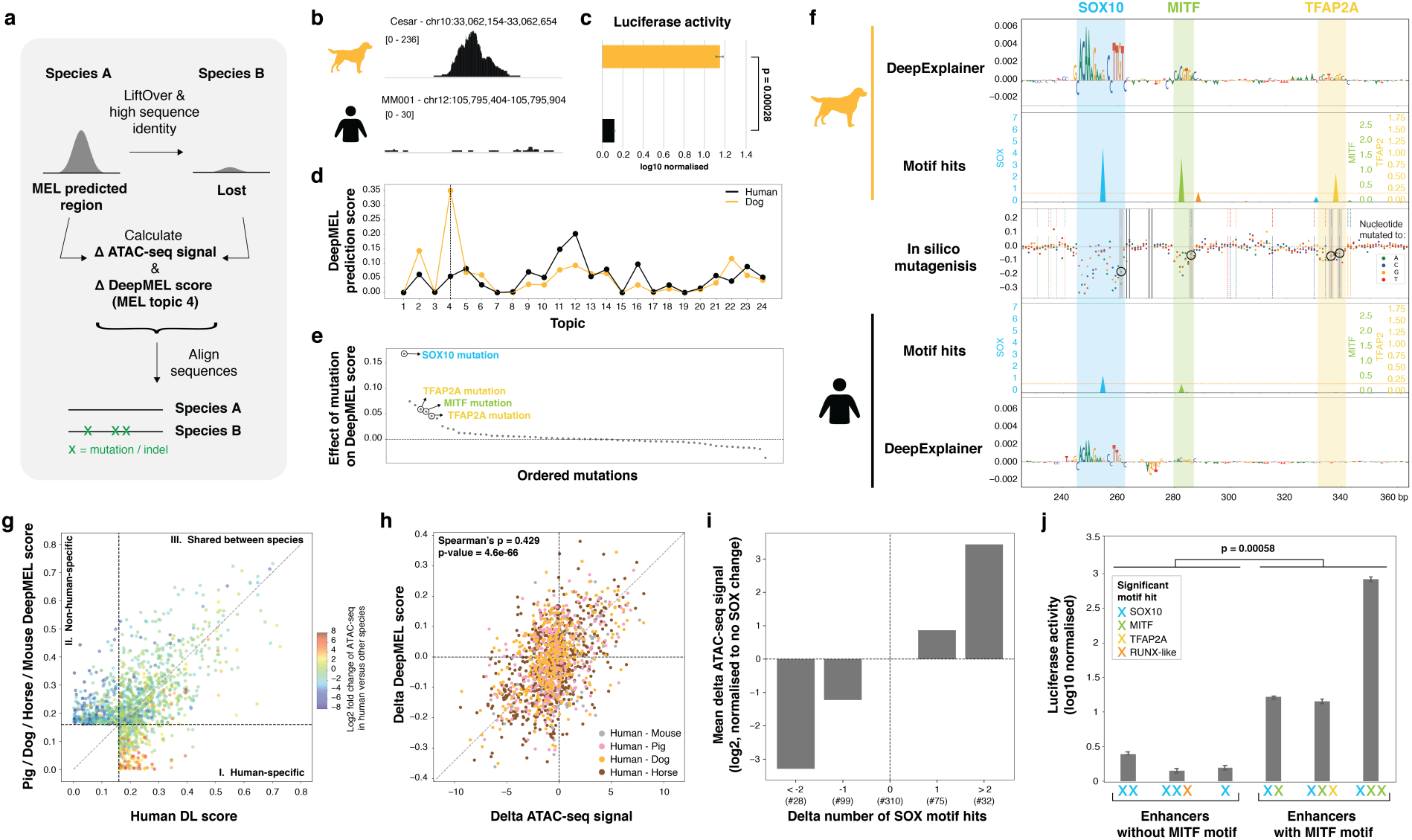
Predicting causal mutations of evolutionary changes in MEL enhancers. **a**, Homologous (identified by stringent liftOver and high sequence identity) MEL enhancers that are accessible and predicted as MEL in one species and that lose accessibility in another are used to identify deleterious cis-regulatory mutations by calculating the delta ATAC-seq signal and delta DeepMEL score for the MEL-specific topic (topic 4). **b, c**, Example region upstream of *APPL2* that is (**b**) accessible and active (**c**) in the MEL dog line Cesar but not in human MEL lines (ATAC-seq profiles of Cesar and MM001 shown here). Luciferase activity in MM001 is shown relative to renilla signal and is log10 transformed. P-value was determined using Student’s t-test and the error bars represent the standard deviation. **d**, DeepMEL prediction score of each of the 24 topics for the dog and human sequence. The dog sequence is predicted as MEL enhancer (topic 4 score > 0.16), whereas this is not the case for the human sequence. **e**, The effect on topic 4 DeepMEL score on the dog sequence when *in silico* simulating each of the single detected point mutations between the dog and human sequence. **f**, DeepExplainer plots and motif hits for SOX10, MITF and TFAP2A are shown for part of the 500 bp dog and human sequence. In the middle, the effect of each possible point mutation between the dog and human sequence on the MEL DeepMEL was *in silico* calculated and is represented by coloured dots depending on the nucleotide the original dog nucleotide was *in silico* mutated to. Truly existing point mutations between the dog and human sequence (as observed by alignment of the sequence via Needle) are highlighted by vertical dashed lines (the colour indicates the original dog base (top dashed line) and the human base (bottom dashed line)). Four mutations that decrease the motif score of the SOX10, MITF and TFAP2A motifs are highlighted by a grey box and are encircled. **g**, Scatter plot of the DeepMEL prediction score for topic 4 in human and in another non-human mammalian species of pairs of homologous sequences. Only enhancers predicted as MEL-specific by DeepMEL (topic 4 score > 0.16) in at least one of the species are used here. Enhancers are represented by a dot and are coloured by the log2 fold change in ATAC-seq signal between human and the other species. In the first quadrant (I) enhancers that are predicted as MEL in human but not in the other species are shown; in quadrant (II) MEL enhancers of non-human species that are not predicted as MEL in human; and the third quadrant (III) contains enhancers that are MEL-predicted in both species. **h**, Scatter plot of the delta ATAC-seq signal and delta DeepMEL prediction score for topic 4 of pairs of homologous enhancers between human and another mammalian species. Dots are colored depending on the species the human homolog was compared to. **i**, Barplot showing the mean effect on the log2 delta ATAC-seq signal of a non-human region compared to the human homolog depending on the number of SOX10 motif hits lost or gained. Only regions having no change in the number of significant TFAP, MITF and RUNX motifs hits were used. The y-axis is normalised to the category with no changes in the number of significant SOX10 motif hits. The number of regions in each of the categories is mentioned (#). **j**. Luciferase assay on six human or dog enhancers. Significant motif hits per enhancer are shown with coloured crosses. Luciferase activity in MM001 is shown relative to renilla signal and is log10 transformed. P-values were determined using Student’s t-test and the error bars represent the standard deviation over three biological replicates.

Next, we performed this analysis on a larger scale, to globally study evolutionary changes in accessibility of orthologous MEL enhancers between human and each of the other mammalian species in our cohort. Firstly, we compared the topic 4 DeepMEL score for each pair of orthologous MEL enhancers and observed that regions predicted as MEL in human but not in the other species were indeed more accessible in human (Fig. 7g, I); in contrast, regions that were only predicted as MEL enhancers in a non-human species were lowly accessible in human (Fig. 7g, II). Orthologous regions that were predicted as MEL enhancer in both human and another mammalian species were similarly accessible in both species (Fig. 7g, III). In fact, DeepMEL proved to be a good predictor for evolutionary changes in accessibility, displaying a high correlation between the delta accessibility and the delta MEL DeepMEL score between orthologous regions (Spearman’s correlation of 0.429) (Fig. 7h). Interestingly, we noticed that among the four CoRC factors, mostly the disruption or gain of one or more SOX10 binding sites between orthologous enhancers quantitatively altered the ATAC-seq signal in a negative and positive way, respectively (Fig. 7i, Fig. S7a), indicating that SOX10 mutations are most causal for changes in MEL enhancer accessibility. Indeed, in the example *APPL2* enhancer presented above, a detrimental mutation in the SOX10 binding site had the strongest effect on the MEL DeepMEL score (Fig. 7e,f), and thus likely, the most impact on not only the loss of enhancer *accessibility* in human (Fig. 7b), but also on the loss of enhancer *activity* (Fig. 7c). However, this was not the case for all MEL enhancers. For instance, an intronic enhancer of *KIF1B* was accessible and predicted as MEL in human, but not in dog (Fig. S7b,d). Although the human region was accessible and predicted as MEL, both the dog and the human enhancer showed no strong activity in a luciferase assay (Fig. S7c). A deeper look at the enhancer code revealed that this human enhancer only contained two significant SOX10 binding sites, but none of the other three CoRC players (Fig. S7e,f). Interestingly, by testing the activity of a total of six human or dog MEL-predicted enhancers, we could distinguish two groups: enhancers that were only accessible and showed little activity (n = 3), or enhancers that were both accessible and significantly more active (n = 3) (Fig. 7j). Profiling DeepExplainer and significant motif hits revealed that the enhancers in the latter group all contained at least one significant MITF binding site, while none of the enhancer in the former group did. Although the number of tested enhancers is small, this trend, together with the fact that MEL enhancers containing a MITF binding site showed increased H3K27ac signal (Fig. S5b), indicates that MITF could function as activator in MEL enhancers. Indeed, MITF has been shown to activate genes involved in pigmentation by recruitment of co-factors and chromatin remodelling complexes^72^ and was previously classified as a TF involved in co-factor recruitment and activation based on its motif distribution in nucleosome depleted regions^6^. Importantly, note that SOX10 binding is insufficient but appears necessary for enhancer activity, as mutations in SOX10 binding sites disrupted enhancer activity in the *IRF4* (Fig. 3g).

In conclusion, DeepMEL provides a suitable platform to study the effect of evolutionary mutations on MEL enhancer accessibility and, in some cases, activity across species. Together, these results validate that SOX10 is crucial for enhancer accessibility in MEL enhancers, and necessary but insufficient for MEL enhancer activity, as activity appeared to be mainly dependent on MITF binding.

## Discussion

Here, we present an in-depth study of melanoma enhancer logic, especially in enhancers specific to the MEL state, by exploiting both cross-species data and machine learning. Although the MEL and MES melanoma cell state have been studied extensively on a transcriptomic and epigenomic level, the combinatorial code of binding sites of their regulatory factors in state-specific enhancers has not yet been explored. Understanding the enhancer logic and the mechanism by which TFs bind and direct active enhancers will become increasingly important, as it will be essential for the development of new therapies that either influence cell state-specific enhancer functions; for the use of (synthetic) enhancers in a targeted way, i.e. enhancer therapy^73,74^; or to prioritise non-coding variants in whole genome sequencing studies of personal or cancer genomes (see our companion paper).

Predicting enhancers and determining their functional role within gene regulatory networks has been an active field for years. Classically, ChIP-seq^1^, motif discovery tools^1,8^, genetic screens^13,14^ and comparative genomic studies^10–12^ have proven useful to reach this goal. For instance Villar et al. uncovered enrichment of CEBPA motifs in highly conserved liver enhancers by performing a comparative genomic analysis in 20 mammalian species; and Prescott et al. identified a novel ‘coordinator’ motif predictive of species-biased cranial neural crest enhancers between human and chimp. Despite the well-established power of cross-species approaches, to our knowledge, a large comparative epigenomics study in melanoma has not yet been conducted, although several non-human models are commonly used in melanoma research^34^. These have either been studied on an intra-species level^33,75–80^; in relation to human melanoma at the level of marker genes^30^, morphology and pharmacological sensitivity^32^, transcriptome^81^; or across three species in the context of genomic landscapes^82^. Here, we conducted a comparative epigenomics study in melanoma across six species, allowing us to demonstrate, for the first time, the conservation of not only the MEL cell state (and the MES cell state in dog), but also the conservation of the underlying master regulators, based on enrichment of TF binding sites within differential MEL and MES peaks and within conserved MEL enhancers.

Although their proven advantages, sequence-based comparative approaches have limited power to identify orthologous regulatory regions in distant species, in part because of the rapid evolution of distal enhancers^83,84^. Methods, such as enhancer element locator (EEL), try to tackle this question by aligning TF binding sites to identify conserved enhancer elements^85^, or by calculating the co-occurrence of sequence patterns^61^. However, these methods are either supervised as they require user-provided PWMs^85^ or are difficult to extract the important biologically-relevant features from^61^. In addition, the identification and exact localisation of important (*de novo*) TF binding sites within enhancers is complex as motif discovery tools are often dependent on user-provided databases and motif-specific thresholds. Recently, deep learning approaches, which are commonly used in disciplines such as speech recognition and image analysis, found their way into the regulatory genomics field to overcome these concerns^15^, but have, to our knowledge, not yet been applied to evolutionary enhancer studies. As deep learning models, such as DeepBind, are particularly powerful in learning complex patterns by leveraging large epigenomics datasets, they are well suited to function as *de novo* motif detectors, as well as to uncover more complex sequence features at higher-level layers that capture the internal structure^15,16^. By designing DeepMEL, a multi-class multi-label neural network trained on melanoma-specific human regulatory topics of co-accessible regions, and by using the model interpretation tool DeepExplainer^54,55^, we were able to perform a thorough and unsupervised analysis of important TF binding sites in melanoma enhancers. Specifically, in MEL enhancers, our data suggests conserved co-binding of a Core Regulatory Complex of four main transcription factors, consisting of SOX10, TFAP2A and MITF. DeepMEL also finds motifs for RUNX factors, but their role in the melocyte or melanoma is less clear. Evidence for co-binding of SOX10, MITF, and TFAP2A was previously observed by enrichment of both MITF and TFAP2A motifs in SOX10 ChIP-seq data in melanoma cells^65^. To predict the precise location and the significance of these TF binding motifs, we designed a new motif scoring scheme by multiplying DeepMEL convolution filters with DeepExplainer profiles^54,55^. We observed high flexibility in the organisation of TF binding sites of the CoRC since eight different modalities were found, formed by all permutations of the CoRC factors, with the exception that all MEL enhancers contained at least one SOX10 binding site. MEL enhancers adhere to a ‘mixed modes enhancer’ model, a billboard-like model with mostly high flexibility in the TF motif organisation, except for the ever-present SOX10 binding sites^68^. Other cross-species studies of enhancers have used ChIP-seq against TFs to examine conserved and divergent enhancers^10,86,87^. Here we avoid the necessity of cross-species ChIP-seq data, as we approximate this by combining ATAC-seq and DeepMEL to characterise, in an unsupervised way, the conservation and divergence of enhancers linked to several melanoma master regulators

It is well recognised that distinct functional classes of TFs exit, with respect to enhancer binding. Pioneer TFs, such as OCT4, SOX2, GRHL, and FOXA1, are able to bind nucleosomal DNA, leading to displacement of the nucleosome and facilitating the binding of other TFs to the accessible enhancer^5,7,68^. SOX2, for example, was shown to bind nucleosomal DNA *in vitro* and associate with closed chromatin^88–90^. SOX2 and other SOX factors have a HMG domain that interacts with the minor groove of the DNA, causing the DNA to bend in a 60-70° angle, a property that has been suggested to contribute to the pioneering activity of SOX2, and possibly of other SOXs^91^. There is still some dispute on the pioneering properties of SOX TFs, as another study classified SOXs as ‘migrant TFs’, i.e. non-pioneering TFs that only bind sporadically to (non)-chromatinised DNA^92^. Nonetheless, we find strong evidence for a pioneering function of SOX10 in MEL melanoma cells. Our current and previous study^29^ have shown that knock-down of SOX10 induces closure of SOX10-bound ATAC-seq peaks containing a SOX10 motif. In fact, DeepMEL predicts SOX10 binding sites as essential for MEL enhancer accessibility. SOX10 is known to engage with open chromatin, as 98% of SOX10 ChIP-seq peaks overlap with DNase-seq sites^57^ and, in addition, SOX10 has been shown to physically interact with BRG1, a subunit of the SWI/SNF chromatin remodeling complex, in differentiating melanocytes^93^. Altogether, this supports the pioneering role of SOX10 in melanocytic melanomas. Notably, especially the binding of SOX10 *dimers* appeared important for MEL enhancer accessibility as eight of the ten enriched SOX10 DL filters in topic 4 represent a SOX10 dimer motif rather than a monomeric motif. This is further supported by the fact SOXE proteins, such as SOX10, are known to form homo- and heterodimers with other SOXE factors^94^. In addition, a study on SOX9, another member of the SOXE TF family, showed that dimerisation of SOX9 was necessary to remodel the chromatin of a *Col2a1* enhancer and to, eventually, allow its activation^95^. Interestingly, we also detected a positional specificity for the SOX10 dimer binding sites as they are mainly localised within the nucleosomal DNA, around 40 bp inwards from the nucleosome start point. Although the findings from Zhu et al. support the binding of SOX(10) proteins inside the nucleosome borders, they observe an enrichment of SOX10 binding towards the dyad of the nucleosome, more towards the center compared to our results reveal. Therefore, further investigations of SOX10 binding to chromatinised DNA might improve the resolution of the exact location of this TF with relation to the nucleosome start and middle point.

Next to pioneer factors, other functional classes of TFs exist, including factors that stabilise the accessibility of the nucleosome depleted regions. TFAP2A was previously classified as such a chromatin stabiliser^6^. Indeed, evolutionary divergence from the TFAP2A consensus motif correlates with loss of chromatin accessibility and H3K27ac ChIP-seq signal^11^. These reports support our observations of TFAP2A as a stabiliser of SOX10-dependent accessible MEL enhancers, likely due to direct competition of TFAP2A with the nucleosome, as TFAP2A binding sites were highly enriched at the predicted center of the central nucleosome. The dependence of SOX10 for opening MEL enhancers prior to TFAP2A binding is in line with the reported classification of TFAP2A as a ‘settler’, a TF whose binding depends predominantly on the accessibility of the chromatin at their binding sites^92^.

Besides classifying accessible (orthologous) regions and predicting important TF motifs within them, DeepMEL is an accurate predictor of the effect of mutations on enhancer accessibility and, for some enhancers, also the activity. This was for instance the case for the *IRF4* MEL enhancer, where DeepMEL performed best among the computational methods tested in Kircher et al.. Note however, that the other models in the benchmark were trained to predict the activity of a total of 20 regulatory regions ranging across different cell types; whereas our DL model is specialised for melanoma regulatory regions. This demonstrates the value of using case-specific training data, such as the data set generated in this study for melanoma. Interestingly, not all predicted MEL enhancers were in fact active. Luciferase assays on a total of six MEL enhancers suggest that SOX10 alone is sufficient for enhancer accessibility, but not for enhancer activity, as MITF binding seems to be needed to activate SOX10-dependent melanoma enhancers. The study of Fufa et al. supports this hypothesis, as activating SOX10-regions in mouse melanocytes showed significant enrichment of E-box motifs (bound by the bHLH protein family, which includes MITF), indicating that it might cooperate with SOX10 to execute melanocyte-specific gene activation. In addition, MITF was previously classified as a TF involved in co-factor recruitment and activation^6,72^. Although SOX10 binding is not sufficient for enhancer activity, it is necessary, as disruption of the SOX10 binding site in the *IRF4* enhancer had a strong effect on activity, probably due to the reappearance of the central nucleosome. Also in an enhancer located about 15 kb upstream of the MEL-specific gene tyrosinase in mouse, both Sox10 and Mitf binding sites were required for activity^96^. This mode of action is also present in other cell types, such as epithelial cells in Drosophila, where Grainyhead acts as pioneer TF and is necessary for both accessibility and activity of epithelial enhancers, but not sufficient for their activity; where it was suggested that the TF Atonal, also a bHLH factor like as MITF, could function as activator of Grh-dependent enhancers^7^. Note that the human and pig predicted MEL enhancers were also accessible in human and pig melanocytes, respectively, indicating that we possibly could extend these observations on the MEL enhancer logic to enhancers in melanocytes.

In conclusion, the combination of comparative epigenomics with deep learning allowed us to perform an in-depth analysis of the melanoma enhancer logic. This work presents an overall framework which can be applied to decipher the enhancer logic in a cell type or cell state of interest, starting from the generation of an extensive cell type-specific (cross-species) epigenomics dataset, all the way through the training and exploitation of a deep neural network to decode enhancer features across species, and to utilise it to assess the impact of cis-regulatory variation.

## Methods

### Cell culture

#### Human melanoma cell lines

Human melanoma cultures (“MM lines”) are short-term cultures derived from patient biopsies^27,35^ (Gembarska et al., 2012; Verfaillie et al., 2015). Cells were cultured at 37°C with 5% CO_2_ and were maintained in Ham’s F10 nutrient mix (Thermo Fisher Scientific) supplemented with 10% fetal bovine serum (FBS; Invitrogen) and 100 µg ml^-1^ penicillin/streptomycin (Thermo Fisher Scientific).

#### Zebrafish melanoma cell lines

Experiments were performed as outlined by Ceol et al.^97^. Briefly, 25 pg of MCR:EGFP were microinjected together with 25 pg of Tol2 transposase mRNA into one-cell Tg(BRAFV600E);p53-/-; mitf-/-zebrafish embryos. Embryos were scored for melanocyte rescue at 48-72 hours post-fertilisation, and equal numbers were raised to adulthood (15-20 zebrafish per tank), and scored weekly (from 8-12 weeks post-fertilization) or bi-weekly (> 12 weeks post-fertilization) for the emergence of raised melanoma lesions^31^. For *in vitro* culture, large tumors were isolated from MCR/MCR:EGFP (14-28 weeks post-fertilization). Zebrafish were maintained under IACUC-approved conditions. Zebrafish primary melanoma ZMEL1 cell line was previously described^38,39^ and EGFP 121-1, EGFP 121-2, EGFP 121-3, EGFP 121-5, were generated as described^98,99^. All cell lines were cultured in DMEM medium (Life Technologies) supplemented with 10% heat-inactivated FBS (Atlanta Biologicals), 1X GlutaMAX (Life Technologies) and 1% Penicillin-Streptomycin (Life Technologies), at 28°C, 5% CO_2_. Zebrafish melanoma lines were authenticated by qPCR and Western for EGFP transgene expression, and periodically checked for mycoplasma using the Universal Mycoplasma Detection Kit (ATCC).

#### Horse melanoma cell lines

The horse cell lines HoMel-L1 and HoMel-A1 are melanoma cell lines derived from a Lipizzaner stallion and Shagya-Arabian mare respectively and were established in Seltenhammer et al.. Cells were cultured at 37°C with 5% CO_2_ in Roswell Park Memorial Institute (RPMI) medium (Thermo Fisher Scientific) supplemented with 10% fetal bovine serum (FBS; Invitrogen) and 1% penicillin/streptomycin (Thermo Fisher Scientific).

#### Pig melanoma and melanocyte cell lines

Both the immortal line of pigmented melanocytes (PigMel) and the primary melanoma cell line (MeLiM) were previously derived^30,100^. PigMel cells were cultured at 37°C with 10% CO_2_ in MEM medium supplemented with 1X MEM non essential amino acids (Thermo Fisher Scientific), 10mM Na pyruvate, 2mM glutamine, 100U/ml penicilin/streptomycin (Thermo Fisher Scientific), 10% FCS and 3,7g/ml Na bicarbonate. MeLiM cells were cultured in DMEM high glucose (Thermo Fisher Scientific), 10% FCS, Pen/Strep, 5% CO_2_.

#### Dog melanoma cell lines

The dog cell lines Bounty and Cesar were established by Aline Primot^37^, and were derived from an uveal melanoma from a Beagle crossed dog and an oral melanoma from the palate from a Shih-tzu, respectively. Cells were cultured at 37°C with 5% CO_2_ in Ham’s F-12 Nutrient Mixture medium (Thermo Fisher Scientific) supplemented with 10% fetal bovine serum (FBS; Invitrogen) and 1% penicillin/streptomycin (Thermo Fisher Scientific).

#### Mouse melanoma cell lines

The mouse melanoma cell line was generated as described in^36^. Cells were cultured at 37°C with 5% CO_2_ in Dulbecco’s Modified Eagle Medium (DMEM) (Thermo Fisher Scientific) supplemented with 10% fetal bovine serum (FBS; Invitrogen) and 1% penicillin/streptomycin (Thermo Fisher Scientific).

### Knock-down experiments

SOX10, TFAP2A and the control knockdown were performed in MM001 using a SMARTpool of four siRNAs against, respectively, SOX10 (SMARTpool: ON-TARGETplus SOX10 siRNA, number L017192-00-0005, Dharmacon), TFAP2A (SMARTpool: ON-TARGETplus TFAP2A siRNA, number L-006348-02-0005, Dharmacon) and a negative control pool (ON-TARGETplus non-targeting pool, number D-001810-10-05, Dharmacon) at a concentration of 20 nM for SOX10-KD, and 40 nM for TFAP2A-KD and the control using as medium Opti-MEM (Thermo Fisher Scientific) and omitting antibiotics. The cells were incubated for 72 h before processing.

### OmniATAC-seq data generation, data processing and follow-up analyses

#### OmniATAC-seq on mammalian lines

OmniATAC-seq was performed as described previously^101^. Cells were washed, trypsinised, spun down at 1000 RPM for 5 min, medium was removed and the cells were resuspended in 1 mL medium. Cells were counted and experiments were only continued when a viability of above 90% was observed. 50,000 cells were pelleted at 500 RCF at 4°C for 5 min, medium was carefully aspirated and the cells were washed and lysed using 50 uL of cold ATAC-Resupension Buffer (RSB) (see Corces et al. for composition) containing 0.1% NP40, 0.1% Tween-20 and 0.01% digitonin by pipetting up and down three times and incubating the cells on ice for 3 min. 1 mL of cold ATAC-RSB containing 0.1% Tween-20 was added and the eppendorf was inverted three times. Nuclei were pelleted at 500 RCF for 10 min at 4°C, the supernatant was carefully removed and nuclei were resuspended in 50 uL of transposition mixture (25 uL 2x TD buffer (see Corces et al. for composition), 2.5 uL transposase (100 nM), 16.5 uL DPBS, 0.5 uL 1% digitonin, 0.5 uL 10% Tween-20, 5 uL H2O) by pipetting six times up and down, followed by 30 minutes incubation at 37°C at 1000 RPM mixing rate. After MinElute clean-up and elution in 21 uL elution buffer, the transposed fragments were pre-amplified with Nextera primers by mixing 20 uL of transposed sample, 2.5 uL of both forward and reverse primers (25 uM) and 25 uL of 2x NEBNext Master Mix (program: 72°C for 5 min, 98°C for 30 sec and 5 cycles of [98°C for 10 sec, 63 °C for 30 sec, 72°C for 1 min] and hold at 4°C). To determine the required number of additional PCR cycles, a qPCR was performed (see Buenrostro et al.^3^ for the determination of the number of extra cycles). The final amplification was done with the additional number of cycles, samples were cleaned-up by MinElute and libraries were prepped using the KAPA Library Quantification Kit as previously described^101^. Samples were sequenced on a HiSeq4000 or NextSeq500 High Output chip.

#### ATAC-seq on zebrafish lines

50,000 cells per line were lysed and subjected to a tagmentation reaction and library construction as described in Buenrostro et al.^3^. Libraries were run on an Illumina HiSeq 2000.

#### Data processing of human melanoma baseline OmniATAC-seq samples

Paired-end reads were mapped to the human genome (hg19-Gencode v18) using bowtie2 (v2.2.6). Mapped reads were sorted using SAMtools (v1.8) and duplicates were removed using Picard MarkDuplicates (v1.134). Reads were filtered by removing mitochondrial reads and filtering for Q>30 using SAMtools. Bam files of technical replicates of the same cell line were merged at this point using samtools merge. Peaks were called using MACS2 (v2.1.2) callpeak using the parameters -q 0.05, --nomodel, --call-summits, --shift -75 --keep-dup all and --extsize 150 per sample. Blacklisted regions (ENCODE) and peaks overlapping with alternative chromosomes and chrM were removed. Summits were extended by 250bp up- and downstream using slopBed (bedtools; v2.28.0), providing human chromosome sizes. Peaks were normalised for the library size using a custom script and overlapping peaks were filtered using the peak score by keeping the peak with the highest score. For visualisation in IGV, normalised bigWigs were made by bamCoverage (Deeptools, v3.3.1), using as parameters --normalizeUsing None, -bl EncodeBlackListedRegions --effectiveGenomeSize 2913022398 and as scaling parameter (-scaleFactor) 1/(RIP/1E6), where the RIP stands for the number of reads in peaks.

#### Data processing of non-human (Omni)ATAC-seq samples, and of human SOX10 and TFAP2A knock-down OmniATAC-seq data

Adapter sequences were trimmed from the fastq files using fastq-mcf (as part of eautils; v1.05) and the read quality was checked using FastQC (v0.11.8). Reads were mapped using STAR (v2.5.1b) (for the zebrafish samples paired-end reads were mapped) to the genome which were downloaded from UCSC (http://hgdownload.cse.ucsc.edu/goldenPath/) (for human: hg19-Gencode v18; for dog: canFam3; for horse: equCab2; for pig: susScr11; for mouse: mm10; for zebrafish: danRer10) and by applying the parameters --alignIntronMax 1 and --aslignIntronMin 2. Mapped reads were filtered for quality using SAMtools (v1.2) view with parameter –q4, sorted with SAMtools sort and indexed using SAMtools index. Peaks were called using MACS2 (v2.1.2) callpeak using the parameters -q 0.05, --nomodel, --call-summits, --shift -75 --keep-dup all and with the genome size for the correct species in --g, and this for each sample per species separately. Summits were extended by 250bp up- and downstream using slopBed (bedtools; v2.28.0), providing the chromosome sizes for the specific species. Per sample, peaks were normalised for the library size using a custom script and overlapping peaks were filtered using the peak score (keeping the highest scoring peak). Normalised bedGraphs were produced by genomeCoverageBed (as part of bedtools; v2.28.0) using as scaling parameter (-scale) 1E6/(number of non-mitochondrial mapping reads). BedGraphs were converted to bigWigs by the bedtools suit functions bedSort to sort the bedGraphs, followed by bedGraphToBigWig to create the bigWigs, which were used in IGV for visualisation.

#### Homer on human and dog differential accessible peaks

First, merged bed files of human and dog ATAC-seq regions were converted to gff format. Count matrices were produced by featureCounts (v1.6.5) using these gff files and bam files of 5 MEL and 5 MES lines for human, and gff and bam files of Cesar and Bounty for dog. Differential peaks were identified using DESeq2 (v1.22.2, R v3.5.2) with a log2FC higher than 2 and a pAdj lower than 0.0005. Homer^47^ was performed on the differential regions using findMotifsGenome.pl, providing the differential regions as a bed file and a fasta file of the human or dog genome, with parameters -mask, - size give and -len 6,8,10,11,12,17,18.

#### Defining sets of conserved ATAC-seq regions

Accessible regions of non-human species were converted to hg19 coordinates using liftOver (Kent-tools) by providing the appropriate liftOver chain (UCSC) and allowing a -minMatch=0.1. LiftOvered regions were intersected with accessible peaks in human (accessible peaks of 5 MEL MM lines) using intersectBed (bedtools, v2.28.0) with -f 0.6 and to define set of conserved regions across species, e.g. conserved regions in across the six species were identified by the intersection of all liftOver bedfiles of non-human species with the human accessible regions, maintaining only the coordinates with which all six species overlapped.

#### Clustering of species based on conserved ATAC-seq regions

Per species, a count matrix was made on the conserved ATAC-seq regions (conserved in all mammalian species or in all six species, as described above) by featureCounts (v1.6.5) using a gff file of the conserved regions in the coordinates of the specific species and bam files for the specific species. Count matrix of different species were merged and the final count matrix was CPM normalised (edgeR v3.22.5, R v3.5.2), followed by quantile normalisation. A principal component analysis (PCA) on the normalised count matrix was performed using irlba (v2.3.3, R v3.5.2) and the first two principal components were used for visualisation.

### Branch length scoring across species

Conserved ATAC-seq regions were identified as described above, and for each of the species, the set of conserved regions was converted to the coordinate system per species and fasta sequences were retrieved. All sequences were scored with our collection of 20,003 motifs using Cluster-Buster^102^ with parameters -m 0, -c 0 and -r 10000. For each motif, the highest CRM score per conserved sequence was used to calculate the BLS across species according to (ref). The branch length was taken from the phylogenetic data from http://hgdownload.cse.ucsc.edu/goldenpath/hg19/phyloP100way/ (UCSC). The sum of the BLSs for all the conserved sequences across the mammalian or all six species was used as a total score for each motif. We normalised these scores by performing BLS on a shuffled variant of all sequences by shuffleseq (EMBOSS, v6.6.0.0), keeping the same base-pair compositions and sequence lengths, and subtracting the shuffled BLS from the true BLS pre motif. This corrected BLS per motif represents the conservation of the motif across a set of conserved regions across a set of species.

### cisTopic analysis to obtain sets of co-accessible regions in human OmniATAC-seq data

To apply cisTopic^29^, a tool for single-cell ATAC-seq analysis, we first simulated single cells form our bulk OmniATAC-seq data on the 17 human melanoma lines via bootstrapping. Per cell line, 50 simulated single cell bam files were generated containing each 50,000 random reads that were bootstrapped from the bulk bam files. These simulated single cell bam files were provided as input for cisTopic (v0.2.0, R v3.4.1), together with the merged regions across all 17 samples, after removing blacklisted regions (ENCODE). We ran cisTopic (parameters: α = 50/T, β = 0.1, burn-in iterations = 500, recording iterations = 1,000) for models with a number of topics between 2 and 30 (2 by 2). The best model, containing 24 topics, was selected on the basis of the highest log-likelihood. Topics were binarised using a probability threshold of 0.995, and performed motif enrichment analysis with cisTarget^8^.

### Deep Learning

#### Data preparation

Regions, which were obtained after peak calling for each baseline (as explained in *Data processing of human melanoma baseline OmniATAC-seq samples*), were merged into one bed file and overlapping regions were removed via custom script. Before intersecting this merged peak file with topics to label each region, regions were augmented in order to have more training data for DeepMEL by extending them to 700 bp and sliding a 500 bp window over them with a 10 bp stride, which meant that each 500 bp augmented region still contained the ATAC-seq summit. Each augmented region had at least 400 bp overlap with its origin. This augmented master region file was intersected with each topic file separately via bedtools and each region was labelled with the topic number if there was an at least 60% overlap. If regions overlapped with multiple topics, we assigned multiple labels to them, allowing for a multi-label and multi-class DL model. The average number of regions in each topic was 1,498 (35,940 in total). After the augmentation and intersection, there were 696,654 regions for training in total, excluding 58,086 chr2 regions for testing.

#### The DeepMEL model architecture and training parameters

The DeepMEL architecture was built by using mainly 4 layers between input and output layer; Conv1D layer (with 128 filters, kernel_size as 20, strides as 1, and activation as relu), MaxPooling1D layer (with pool_size as 10 and strides as 10), TimeDistributed Dense layer together with Bidirectional LSTM layer (with 128 units, dropout as 0.1, recurrent_dropout as 0.1), and Dense layer (with 256 units and activation as relu). After MaxPooling1D, Bidirectional LSTM, and Dense layer Dropout was used as 0.2, 0.2, and 0.4 respectively. The DeepMEL takes one-hot encoded (500bp x 4 nucleotide) forward and reverse strand of the region, passes them separately through the model and takes the average of the activations of the neurons in the final Dense layer (24 units corresponding to 24 topics with sigmoid activation) with the *average* function in order to make the final prediction. The model was compiled using Adam optimizer with 0.001 learning rate. In order to make the model multi-label classifier, sigmoid activation function was used at the end of the final layer of the model and binary cross entropy loss function was used. The model was trained for 2 epochs with 128 batch size, which took 67 minutes. Keras 2.2.4^103^ with tensorflow 1.14.0^104^ was used. A Tesla P100-SXM2-16GB GPU was used for training on VSC servers (Flemish Supercomputer Center).

#### Performance evaluation

The performance of the model was evaluated for each topic separately since it was a multi-label classifier. The area under the Receiver Operator Characteristic curve (auROC) and the Precision Recall curve (auPR) were calculated for training (regions on all chromosomes except chr2), test (regions on chr2), and label-shuffled regions.

#### Converting convolution filters to PWMs, filter-topic assignment, and filter-annotation

After the model was trained, the filters of the convolution layer were converted into PWMs by the following strategy: (i) 4,000,000 unique 20bp-long (size of the filters) sequences were randomly generated. (ii) The activation score of each filter for each sequence was calculated and the top 100 sequence were selected. (iii) A count matrix was generated from these 100 sequences obtained for each filter. (iv) Finally, the count matrices were converted into PWMs. In order to assign the filters to topics, a similar strategy that is mentioned in Basset^18^ was used. The activation score of the filter was separately set to its mean activation score over all sequences, then the loss/accuracy score on the prediction was calculated for each class. Filters were ordered based on their effect on a certain topic. After the filters were converted into PWMs, Tomtom^59^ motif annotation tool was used together with using a curated collection of more than 22,000 PWMs in order to annotate the DL features to known motifs. The cutoff for the q-value was set to 0.3.

#### DeepExplainer

Among 35,940 topic regions, 500 of them were randomly selected to initialise DeepExplainer^54^. Importance score for each position of the sequence of interest was calculated with respect to any of the 24 classes. The hypothetical importance score, which is obtained from the DeepExplainer output, was multiplied by the one-hot encoded matrix of the sequence. Finally, the 500 bp sequences were visualised by adjusting the nucleotide heights based on their importance score by using modified viz_sequence function from the DeepLift^105^ repository.

#### In silico saturation mutagenesis on IRF4

By changing each nucleotide on a 500 bp sequence into three other nucleotides, 1,500 sequences that contain only one mutation compared to initial sequence were generated and scored by the model. The delta prediction score for each mutation was calculated for each class by comparing the final prediction score relative to the prediction score for the initial sequence. The *IRF4* enhancer (chr6:396,143-396,593) used in *in vitro* saturation mutagenesis assay is also covered by one of our MEL enhancers predicted as topic 4 (chr6:396,135-396,636). *In silico* saturation mutagenesis assay on this region was done using the delta prediction score of topic 4 and a Pearson correlation was calculated on overlapping nucleotides between the *in silico* and *in vitro* assays (451 bp).

#### Motif scoring method and centering regions

Using only the filters identified from the convolutional layer is not sufficient to localise significant motif hits on MEL enhancers since it does not necessarily mean that when the activation of a filter passes the activation threshold, that the filter has an effect on the final classification for a position in the sequence. Also using only the DeepExplainer importance scores is not sufficient either since it is not able to precisely detect the exact location, size, and the name of the motif hit. In order to overcome this problem, activation scores of the filters on each sequence were multiplied by the DeepExplainer importance scores. Then, a threshold was calculated for each motif by comparing MEL and MES enhancers after the output of the multiplication was normalised. This approach yielded significant motif hits with their precise location.

#### Nucleosome positioning

Nucleosome start and middle point predictions were calculated by using an executable nucleosome prediction tool called Kaplan_v3^69^ that takes only the DNA sequence and calculates the nucleosome positioning for each nucleotide. In order to get more precise results, as the authors of Kaplan_v3 suggest, enhancers were extended 3 kb from both ends. After obtaining the predictions, the middle 500 bp part of the 6.5kb nucleosome prediction score was used.

#### Tn5 footprinting

Footprint of the Tn5 was determined by inferring Tn5 cut sites with a custom script that takes bam file sand locates the Tn5 cut site deduced from the start point of each read resulted from the ATAC sequencing.

### AUROC on human and dog of DL and Cluster-Buster

To the performance of the model to discriminate between MEL and MES regions in human and dog was performed by scoring the top 5,000 differential MEL and MES regions in human and dog (described above) by DeepMEL and calculating precision of correct assignment (i.e. topic 4 score for the MEL regions and topic 7 scores for the MES regions). The performance of DeepMEL was compared with the motif scoring tool Cluster-Buster^102^ by scoring the same sets of regions with Cluster-Buster by using a merged motif file of (some of) the top filters identified by the model in either topic 4 or topic 7, and by using the obtained CRM score to estimate the performance of Cluster-Buster.

### Identification of homologous MEL genes and enhancers

To identify genes differentially expressed in human MEL cell lines, we performed DEseq2 (v1.22.2, R v3.5.2) on 7 MEL (MM031, MM034, MM057, MM074, MM087, MM118, MM164) and 5 MES (MM029, MM099, MM116, MM163, MM165) human lines. 379 genes were found differentially expressed in MEL lines (log2FC > 2.5 and adjP < 0.005). We converted the gene symbols to Ensemble gene IDs using biomaRt (v2.38.0, R v3.5.2) and found back the genomic locations of the genes using GenomicFeatures (v1.34.8, R v3.5.2). We searched for MEL enhancers in the extended gene loci, by extending the genomic locations 200 kbp upstream and downstream of the start and the end of the gene and using bedtools intersect (v2.28.0) to intersect the extended loci with the MEL-predicted regions in MM001. For the human differential MEL genes with at least one MEL-predicted peak in their extended gene locus, the homologous genes in the other six species was identified by using biomaRt to convert the human Ensemble gene IDs to Ensemble gene IDs of the other species. Again GenomicFeatures was used to get the genomic locations of the genes in the different species. Next, we identified the MEL enhancers per species that were intersection with the extended gene loci of each of the homologous genes in that specific species using bedtools intersect. liftOver -minMatch=0.1 was used to calculate the number of these regions that could be identified by performing coordinate conversion.

### Correlation of MEL enhancers using deep layers of DeepMEL

Conserved MEL enhancers in the extended loci of conserved MEL genes across the six species were scored by the DeepMEL. By taking the activation scores of the neurons on the Dense layer, which comes before the final output layer and harbours the characteristics and the contents of the enhancers coming from previous feature extraction layers, a matrix was generated consisting of a score for 256 nodes for each of the regions. A pearson correlation was generated to calculate the pairwise similarity between each of the regions.

### Genome-wide prediction of MEL enhancers

(Soft)-masked genomes where downloaded from UCSC for Homo sapiens (human, hg19), Equus caballus (horse, equCab2), Sus scrofa (pig, susScr11), Canis lupus familiaris (dog, canFam3), Mus musculus (mouse, mm10), Danio rerio (zebrafish, danRer10), Ciona intestinalis (ci3), Caenorhabditis elegans (ce11) and Saccharomyces cerevisiae (sacCer3). The first chromosome of each species was tiled with a sliding window of 500 bp and a 100 bp shift using bedtools makewindows (v2.28.0). Tiles containing ‘N’ were deleted and the remaining tiles were scored by DeepMEL. The number of MEL-predicted tiles (topic 4 score > 0.16) was divided by the number of genes per species to yield an estimate of the content of the MEL-enhancer code in each genome.

### Mutations in orthologous enhancers across species

We defined highly-probable orthologous MEL enhancers between human and another species as regions that were predicted as MEL in one species and for which there was a stringent liftOver (liftOver -minMatch=0.995) and high sequence identity (more than 80% after pairwise alignment via needle (EMBOSS, v6.6.0.0), using parameters -gapopen 10.0 -gapextend 0.5) in the other species. Note that also the reverse complement of the regions was checked here. Delta ATAC-seq scores were calculated for the pairs of orthologous regions by making a count matrix using featureCounts (v1.6.5) on the regions and the bam file of a sample of the species, and by normalising this count matrix using the library size according to the bam file used, followed by dividing the counts of the two species (human counts / non-human counts) after adding a pseudocount. Mutations were identified by alignment via needle, using parameters -gapopen 10.0 -gapextend 0.5.

### Luciferase assay

Six MEL-predicted enhancers (3 in the dog line Cesar and 3 in the human line MM001) were synthetically generated and cloned into a pTwist ENTR plasmid (Twist Bioscience) via Twist Bioscience. Regions were transferred from the Gateway entry clone into the destination vector (pGL4.23-GW, Addgene) via an LR reaction by mixing 2 ul of the entry clones (100 ng/ul) with 1ul of the destination plasmid (150 ng/ul), 1 ul TE buffer and 1 ul LR enzyme (LR Clonase II Plus enzyme mix, Thermo Fisher Scientific), and incubation at 25°C for 1 hour. Afterwards, 1ul of proteinase K (Thermo Fisher Scientific) was added and reactions were incubated at 25°C for 10 min. 3ul of each LR reaction was transformed into 50 ul of Stellar competent cells (Takara Bio) via heatshock, 200ul of SOC medium was added and incubated for 1 hour in a shake incubator at 37°C, before plating the transformed cells on LB agar plates with 1/1000 carbenicillin and incubation overnight at 37°C. One colony per construct was grown overnight in a shake incubator at 37°C before plasmid extraction using the NucleoSpin Plasmid Transfection-grade kit (Macherey-Nagel). For each construct three biological replicates were performed by transfecting the plasmids into 80% confluent cells of MM001 in a 24 well plate. Per transfection, 400ng of the construct was transfected together with 40ng of Renilla plasmid (Promega) using lipofectamine 2000 (Thermo Fisher Scientific). Luciferase activity of each construct was measured using the Dual-Luciferase Reporter Assay (Promega) according to the manufacturer’s instructions. Luciferase activity was normalised against the Renilla luciferase activity.

### Publicly available data used in this work

SOX10 ChIP-seq and MITF ChIP-seq data on the 501Mel melanoma cell lines were downloaded as raw fastq files from NCBI’s Gene Expression Omnibus through GEO accession number GSE61965 ^65^ and were mapped to the human genome using Bowtie2 (v2.1.0) and peaks were called by MACS2 (v2.1.1). TFAP2A ChIP-seq data on human primary melanocytes from neonatal foreskin was retrieved from Seberg et al. (GSE67555) as a bed file, which was converted to a bedGraph and BigWig using the peak height from the bed file. H3K27ac-seq and H3K27me3 ChIP-seq data for MM001 (GSE60666); and RNA-seq data (data for MM031, MM034, MM057, MM074, MM087, MM099 and MM118 was downloaded from GSE60666; data for MM029, MM116, MM0163, MM164, adn MM165 from GSE134432) were processed as mentioned in Verfaillie et al.. OmniATAC-seq data for the human lines MM001, MM011, MM029, MM031, MM047, MM074, MM057, MM087 and MM099 were obtained through GSE134432^28^ and were processed as described above in ‘Data processing human melanoma baseline OmniATAC-seq samples’; which was also the case for ATAC-seq data from normal human melanocytes on foreskin (NHM1), which were downloaded as raw fastq files from GSE94488 (GSM2476338)^106^. ATAC-seq data from C. elegans and S. cerevisiae were downloaded as raw fastq files from GSE114439 (SRR7164221)^107^ and GSE66386 (SRR1822137)^108^, respectively, and were mapped paired-end using STAR (v2.5.1b) to ce11 and sacCer3, respectively, before calling peaks using MACS2 (v2.1.2) with -q 0.05, extending the peaks 250bp up- and downstream of the summit and filtering out overlapping peaks based peak height. The MPRA data on the *IRF4* enhancer was downloaded from https://mpra.gs.washington.edu/satMutMPRA/ and was processed as described above.

### Data availability

The data generated for this study have been deposited in NCBI’s Gene Expression Omnibus and are accessible through GEO Series accession number GSE142238. This includes OmniATAC-seq data of eight human melanoma cell lines, two dog melanoma cell lines, two horse melanoma cell lines, one pig melanoma cell line, one pig melanocyte cell lines and one mouse melanoma cell line; ATAC-seq data of four zebrafish cell lines and OmniATAC-seq data of SOX10 and TFAP2A knock-down in the human melanoma cell line MM001.

## Acknowledgements

This work was supported by funded by an ERC Consolidator Grant to S.A. (no. 724226_cis-CONTROL), by the KU Leuven (grant no. C14/18/092 to S.A.), by the Foundation Against Cancer (grant no, 2016-070 to S.A.), a PhD fellowship from the FWO (L.M., no. 1S03317N) and a postdoctoral research fellowship from Kom op tegen Kanker (Stand up to Cancer), the Flemish Cancer Society, and from Stichting tegen Kanker (Foundation against Cancer), the Belgian Cancer Society (J.W.). We would like to thank Odessa Van Goethem and Véronique Benne for their contribution in establishing and providing the mouse melanoma cell line, and Leif Andersson for sharing the horse melanoma cell lines. Computing was performed at the Vlaams Supercomputer Center and high-throughput sequencing via the Genomics Core Leuven. The funders had no role in study design, data collection and analysis, decision to publish or preparation of the manuscript.

## Author contributions

L.M., I.I.T. and S.A. conceived the study. L.M. performed the experimental work for the mammalian OmniATAC-seq dataset with the help of L.V.A, S.M., V.C and J.W.. M.F., E.v.R. and L.Z. established and maintained the zebrafish cell lines and performed ATAC-seq on these. G.E.M. maintained and provided the pig cell lines. A.P. and E.C. established and provided the dog cell lines. P.K. established and provided the mouse melanoma cell line. M.S. established, maintained and provided the horse cell lines. G.E.G. estabiled and provided the human cell lines. L.M. performed the experimental work and analysis of the luciferase assays together with D.M. L.M. performed the bioinformatic analyses of the OmniATAC-seq dataset. I.I.T. established the neural network and performed all bioinformatic analyses regarding the model. L.M., I.I.T., J.W. and S.A. wrote the manuscript.

