## Supplementary figures for "Cross-species analysis of melanoma enhancer logic using deep learning"

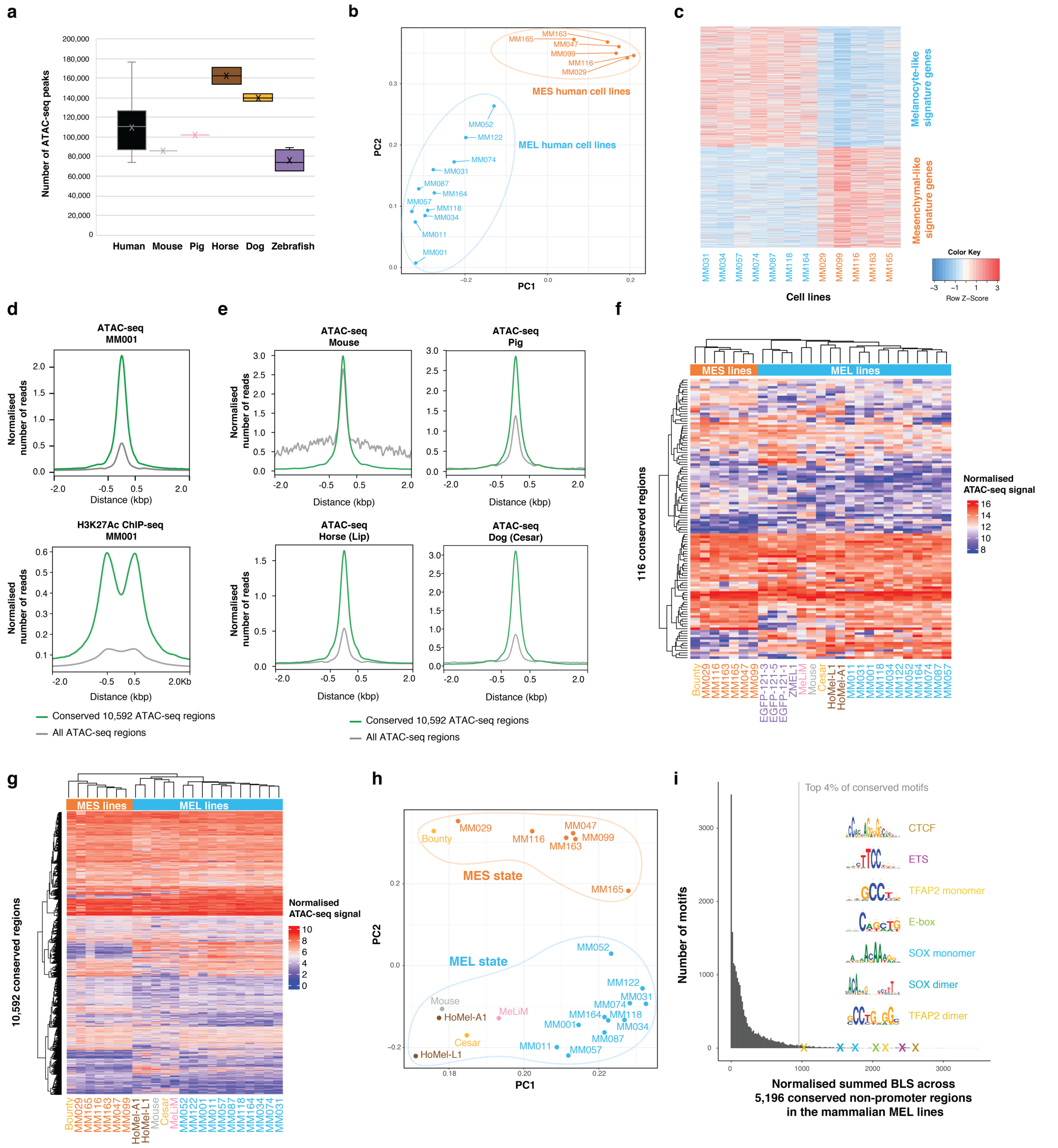

**Figure S1. Clustering of melanoma samples across species**. **a**, Distribution of the number of ATAC-seq peaks called per sample across species. **b**, PCA plot of the human melanoma lines using counts on the 50,000 most variable peaks. Human cell lines cluster into two distinct groups that are linked to the two main melanoma states. **c**, Expression of MEL and MES signature genes on 12 of the human cell lines. **e**, ATAC-seq and H3K27ac ChIP-seq signal of the human line MM001 on the 10,592 conserved regions and on all human regions. **e**, ATAC-seq signal of mouse, pig, horse (Lip) and dog (Cesar) melanoma samples on the 10,592 conserved regions and on all regions of the specific species. **f, g**, Heatmap of ATAC-seq signal of (**f**) all samples on the 116 conserved regions and (**g**) of all mammalian samples on the 10,592 mammalian-conserved regions. **h**, PCA plot of the mammalian melanoma lines based on the 10,592 conserved regions. **g**, Histogram of the cumulative BLS for 20,003 motifs on 5,196 conserved non-promoter regions (i.e. no proximal promoters situated < 1 kbp from TSS) in the MEL lines across the mammalian species. The first hit of the top recurrent TF binding motifs within the top 4% conserved motifs is indicated as a cross and is accompanied by the logo of the motif.

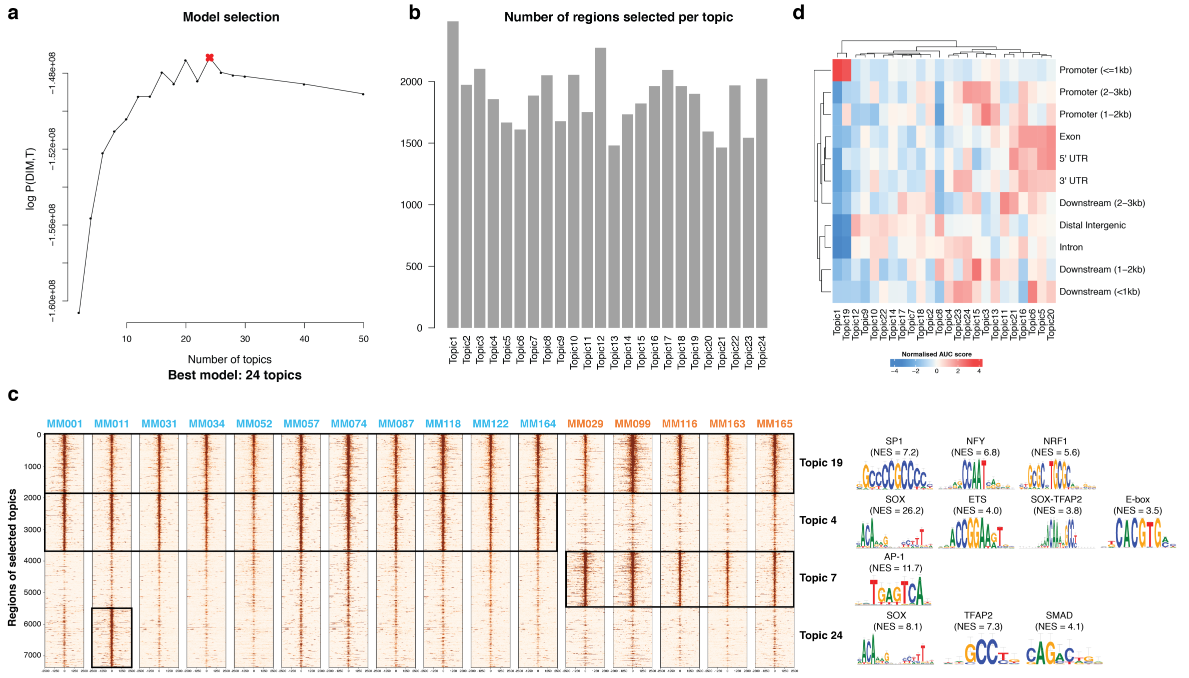


**Figure S2. Topic modeling on human ATAC-seq data**. **a**, cisTopic model with 24 topics was selected based on the log-likelihood in the last iteration. **b**, Number of regions per topic after topic binarisation. **c**, Coverage heatmaps of ATAC-seq data to validate the regions per topic for four selected topics: a promoter topic (topic 19), the MEL-specific topic (topic 4), the MES-specific topic (topic 7) and a cell line-specific topic (topic 24, specific to MM001). Key motifs found enriched in the selected topics by i-cisTarget, accompanied by their NES score. **d**, Topic annotation heatmap.

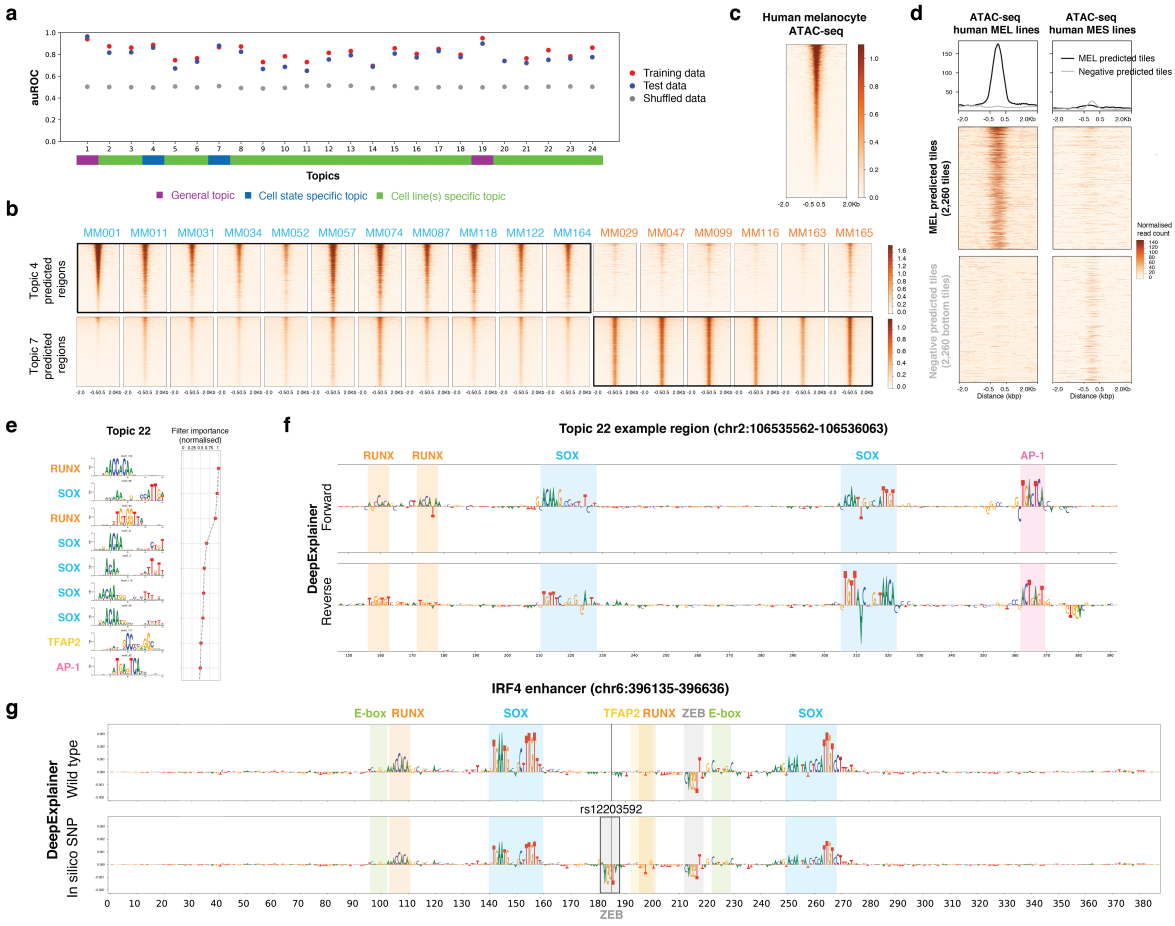


**Figure S3. DeepMEL reveals important features in melanoma regulatory classes. a,** Area under the receiver operating characteristics (auROC) curve of DeepMEL for training, test and shuffled data for classifying regions of the 24 topics. Note that promoter topic regions are the easiest to classify by the model. **b**, Coverage heatmaps of ATAC-seq data of the 17 human lines to validate topic regions predicted by DeepMEL for the MEL-specific topic (topic 4) and MES-specific topic (topic 7). **c**, ATAC-seq signal of melanocytes on topic 4 predicted regions by DeepMEL in MM001, ordered according to the melanocyte ATAC-seq signal. **d**, ATAC-seq signal of merged human MEL lines and merged human MES lines on 2,260 MEL-predicted tiles (DeepMEL score of topic 4 > 0.16) when scoring 500bp sequences tiled across chromosome 1 in the human genome by DeepMEL, and on the 2,260 tiles with the lowest topic 4 prediction score. Heatmaps are coloured by normalised read counts and aggregation plots are shown on top. **e**, The top nine features learned DeepMEL to classify topic 22 regions (regions specific to the intermediate state). **f**, DeepExplainer plots on the forward and reverse strand of an example topic 22 region. The enhancer contains both AP-1 and SOX motifs. **g**, DeepExplainer profiles of wildtype *IRF4* enhancer and after *in silico* introduction of the rs12203592 SNP, which creates a ZEB-like motif.

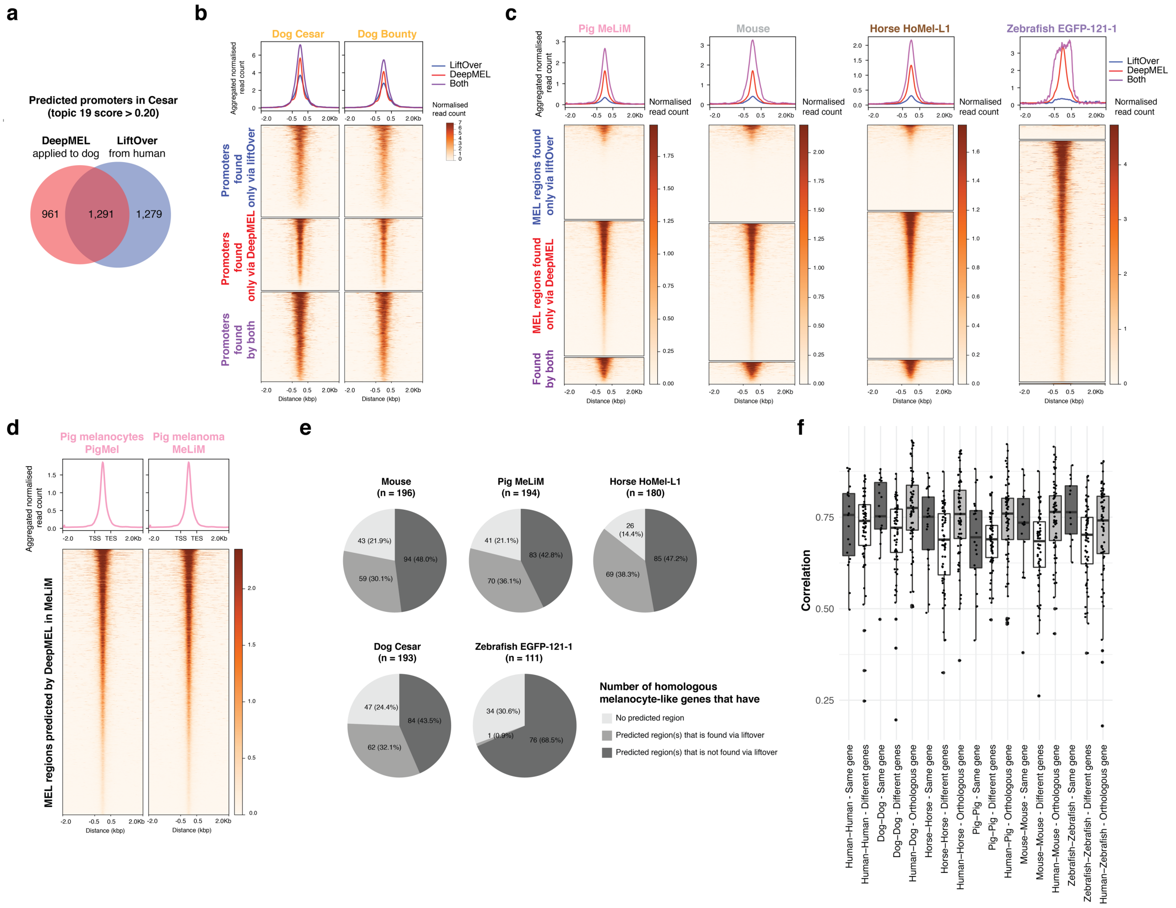

**Figure S4. Genome-wide MEL enhancer predictions using DeepMEL. a**, Venn diagram of the number of topic 19 (i.e. promoter) regions predicted by DeepMEL in dog or found via liftOver of the human topic 19 predicted regions to dog coordinates. **b**, Heatmaps of ATAC-seq signal of the dog lines ‘Cesar’ and ‘Bounty’ on topic 19 regions found via liftOver (blue), topic 19 regions predicted by DeepMEL (red) and topic 19 regions identified by both methods (purple). Heatmaps are coloured by normalised read counts and ordered according to the ATAC-seq signal in ‘Cesar’. Aggregation plots of are shown on top. **c**, Heatmaps of ATAC-seq signal of pig, mouse, horse and zebrafish on MEL regions regions found via liftOver (blue), MEL regions predicted by DeepMEL (red) and MEL regions identified by both methods (purple). Heatmaps are coloured by normalised read counts and ordered according to the ATAC-seq signal in ‘Cesar’. Aggregation plots of are shown on top. **d**, ATAC-seq signal of pig melanocytes (‘PigMel’) and pig melanoma (‘MeLiM’) on MEL regions predicted in ‘MeLiM’ ATAC-seq regions. Heatmaps are coloured by normalised read counts and aggregation plots of are shown on top. **e**, Per species, the number of orthologous of the 217 human MEL-differential genes and that had a topic 4 predicted region in 200 kb up- and downstream of the gene in the respective species are shown between brackets. Of these genes, boxplots show per species the number of genes that do not have a MEL-predicted regions within their 200 kb up- and downstream extended gene locus, that have a MEL-predicted region within their extended locus which was either also found by liftOver or not identified via liftOver. **f**, Pearson correlation of deep layer scores between MEL-predicted regions of homologous genes between human and another species (‘Human-Species - Orthologous gene’), between MEL-predicted regions near different genes within one species (‘Species-Species - Different gene’) and between MEL-predicted regions near the same gene within one species (‘Species-Species - Same gene’).

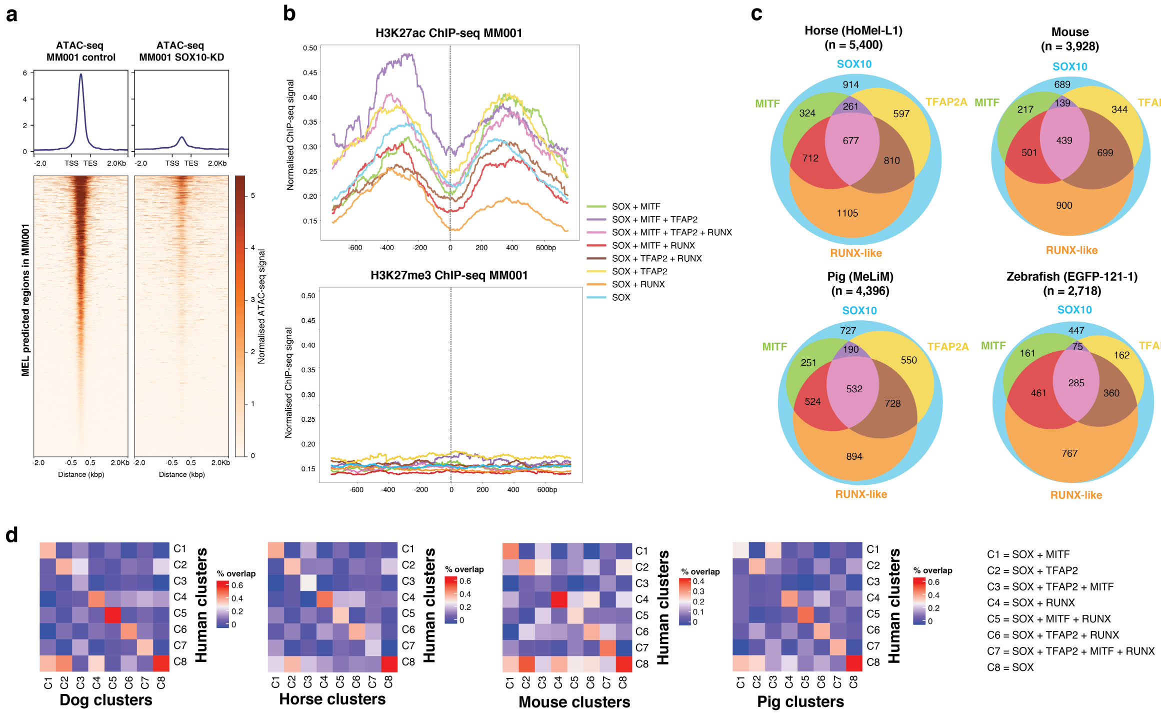


**Figure S5. Conservation of COre Regulatory Complex clusters in MEL enhancers.** **a**, Heatmaps of ATAC-seq signal of MM001 after 72h of control knock-down (DMSO) and SOX10-KD on the 3,885 MEL predicted regions in MM001. Heatmaps are coloured by normalised read counts and ordered according to the ATAC-seq signal in the control sample. Aggregation plots of are shown on top. **b**, Aggregation plots of H3K27ac and H3K27me3 ChIP-seq in MM001 on the human MEL enhancer clusters. **c**, Venn diagrams of the enhancer clusters of DeepMEL-predicted MEL enhancers in horse, mouse, pig and zebrafish. The number of predicted MEL enhancers in each species is mentioned between brackets. **d**, Heatmaps showing correlation between MEL enhancers clusters of human and each of the other mammalian species. Heatmaps are coloured by the percentage of overlap between regions after liftOver of the enhancer coordinates to human.

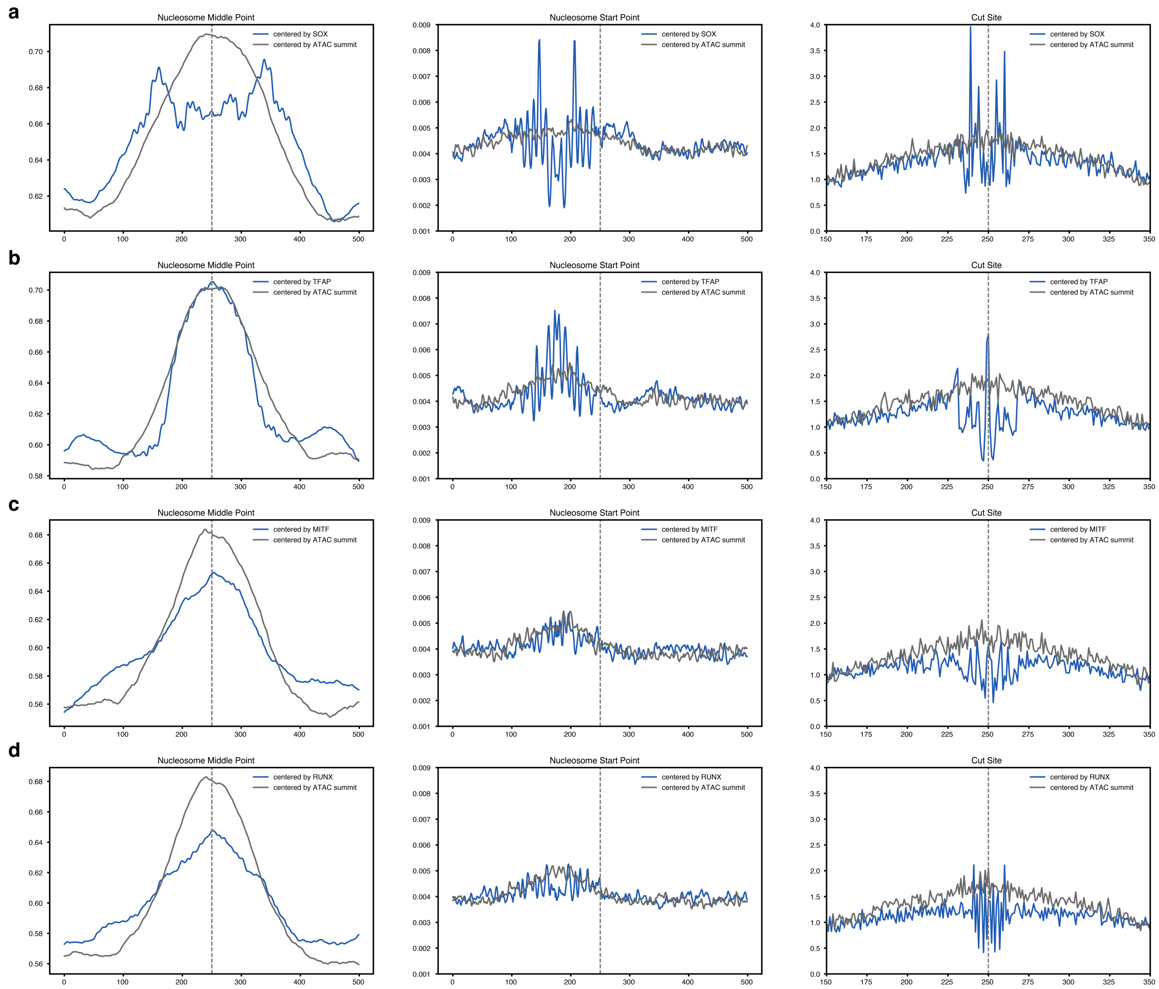


**Figure S6. SOX10, TFAP2A, MITF and RUNX TF binding site motifs in relation to the nucleosome**. **a,b,c,d**, Nucleosome middle point (left), nucleosome start point (middle) and Tn5 cut site (right) on MEL-predicted regions containing one SOX10 motif (**a**), one TFAP2 motif (**b**), one MITF motif (**c**) and one RUNX-like motif (**d**) next to possible other motifs, where the regions are either centered on the ATAC-seq summit (grey) or specified motif (blue).

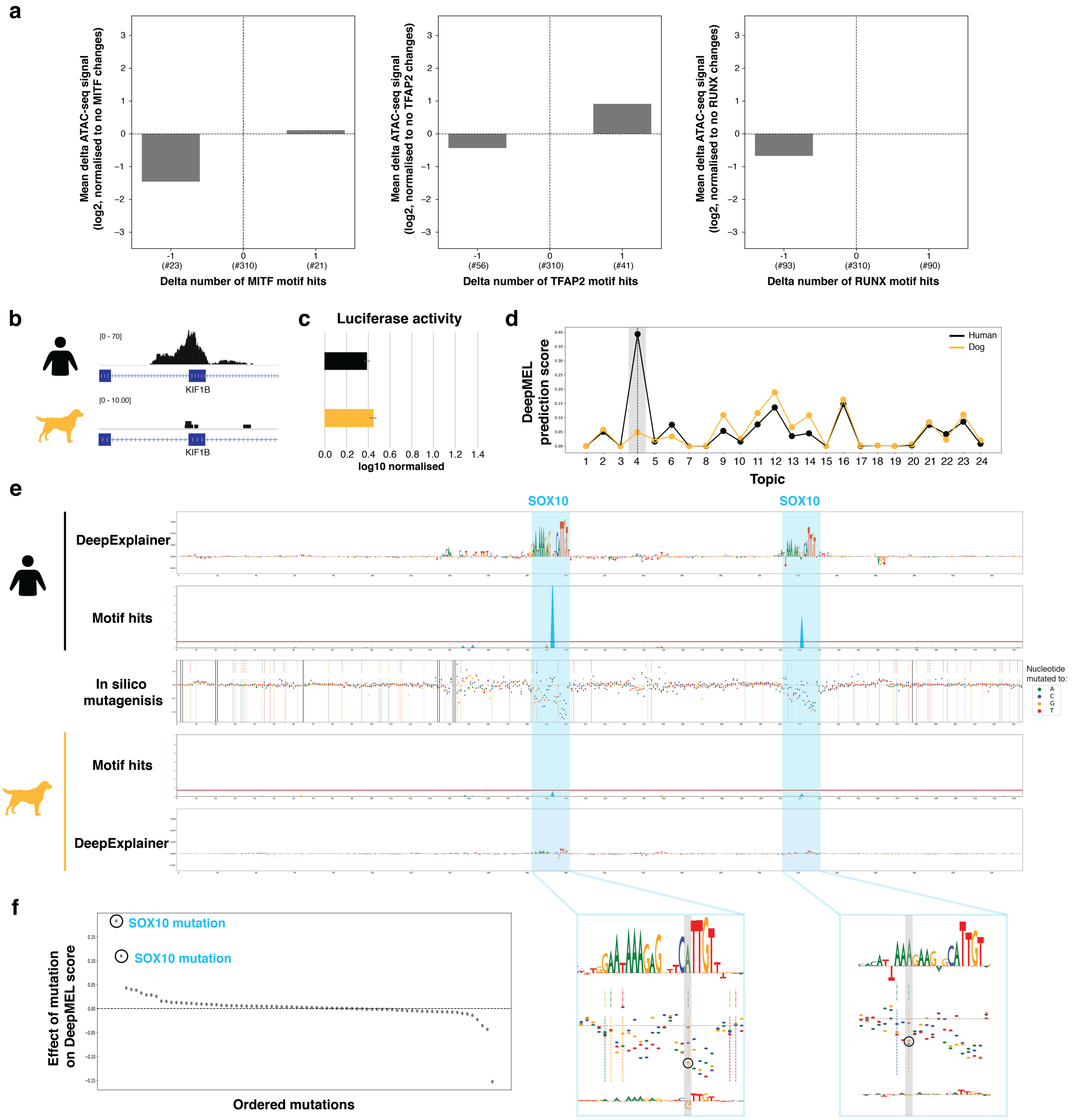


**Figure S7. SOX10 binding site disruption destroys enhancer accessibility in *KIF1B* MEL enhancer. a**, Barplot showing the mean effect on the log2 delta ATAC-seq signal of a non-human region compared to the human homolog depending on the number of MITF, TFAP or RUNX motif hits lost or gained. Only regions having no change in the number of significant motifs hits for the other three CoRC factors were used. The y-axis is normalised to the category no changes in the number of significant motif hits. The number of regions in each of the categories is mentioned (#). **b**, Example MEL enhancer of *KIF1B*, which is accessible in human but not in dog. **c**, Both the human and dog regions show nearly no enhancer activity as measured by a luciferase assay in MM001. **d**, The region is predicted as MEL in human but not in dog. **e**, DeepExplainer profiles for human and dog are shown for the *KIF1B* region together with motifs hits for SOX10, TFAP2A, RUNX and MITF. Only for SOX10, two significant motif hits (peak above the threshold line) are visible in the human sequence. The middle row shows indels (black dashed lines) and individual point mutations (indicated by dashed lines, the colour indicates the human base (top dashed line) and the dog base (bottom dashed line)) and the dots represent the effect on the MEL DeepMEL score of the point mutations. In the zoom-in, the two A-to-G mutations that cause disruption of SOX10 motif hits in the *KIF1B* region in dog are shown. **f**, The effect on the MEL prediction score could mainly be explained by two mutations, which correspond to the point mutations in the two SOX10 motifs.
